# Blocks in tricarboxylic acid cycle of *Salmonella enterica* cause global perturbation of carbon storage, motility and host-pathogen-interaction

**DOI:** 10.1101/832675

**Authors:** Janina Noster, Nicole Hansmeier, Marcus Persicke, Tzu-Chiao Chao, Rainer Kurre, Jasmin Popp, Viktoria Liss, Tatjana Reuter, Michael Hensel

**Affiliations:** Abt. Mikrobiologie, Universität Osnabrück, Osnabrück, Germany; Department of Biology, Luther College at University of Regina, Regina, Canada; Microbial Genomics and Biotechnology, Center for Biotechnology, Universität Bielefeld, Bielefeld, Germany; Institute of Environmental Change & Society, University of Regina, Regina, Canada; iBiOS; CellNanOs, Universität Osnabrück, Osnabrück, Germany

**Keywords:** TCA cycle, glycogen metabolism, chemotaxis, phagocytosis

## Abstract

The tricarboxylic acid cycle is a central metabolic hub in most cells. Virulence functions of bacterial pathogens such as facultative intracellular *Salmonella enterica* serovar Typhimurium (STM) are closely connected to cellular metabolism. During systematic analyses of mutant strains with defects in TCA cycle, a strain deficient in all fumarase isoforms (Δ*fumABC*) elicited a unique metabolic profile. Alongside fumarate STM Δ*fumABC* accumulates intermediates of glycolysis and pentose phosphate pathway. Analyses by metabolomics and proteomics revealed that fumarate accumulation redirects carbon fluxes towards glycogen synthesis due to high (p)ppGpp levels. In addition, we observed reduced abundance of CheY, leading to altered motility and increased phagocytosis of STM by macrophages. Deletion of glycogen synthase restored normal carbon fluxes and phagocytosis, and partially levels of CheY. We propose that utilization of accumulated fumarate as carbon source induces a status similar to exponential to stationary growth phase transition by switching from preferred carbon sources to fumarate, which increases (p)ppGpp levels and thereby glycogen synthesis. Thus, we observed a new form of interplay between metabolism of STM, and cellular functions and virulence.

**Importance:** We performed perturbation analyses of the tricarboxylic acid cycle of the gastrointestinal pathogen *Salmonella enterica* serovar Typhimurium. The defect of fumarase activity led to accumulation of fumarate, but also resulted in a global alteration of carbon fluxes, leading to increased storage of glycogen. Gross alterations were observed in proteome and metabolome compositions of fumarase-deficient *Salmonella*. In turn, these changes were linked to aberrant motility patterns of the mutant strain, and resulted in highly increased phagocytic uptake by macrophages. Our findings indicate that basic cellular functions and specific virulence functions in *Salmonella* critically depend on the proper function of the primary metabolism.

## Introduction

The central carbon metabolism (CCM) is essential for all prototrophic bacteria because it provides energy, as well as precursors for biosynthesis of a large number of biomolecules. In particular, the tricarboxylic acid cycle (TCA cycle) produces the reductive equivalents for the electron transport chain and the carbon backbone for various amino acids, making it an important hub for efficient bacterial metabolism in changing environments (1, 2). Several endogenous factors, such as the energy status of the cell, influence TCA cycle activity. For example, the activity of the isocitrate dehydrogenase is allosterically stimulated by ADP (3), whereas α-ketoglutarate dehydrogenase is inhibited by its products succinyl-CoA and NADH (4). In addition, bacterial citrate synthesis is controlled by allosteric inhibition of citrate synthase by ATP and NADH (5). However, TCA cycle activity is also influenced by exogenous factors, such as exposure to antibiotics and ROS, which target sensitive enzymes harboring Fe-S clusters (6, 7).

*Salmonella enterica* serovar Typhimurium (STM) is an invasive facultative intracellular pathogen, the causative agent of human gastroenteritis, and serves as model organism for systemic *Salmonella* infections. The divergent host niches colonized during infection demand STM to adapt its metabolism from the intestinal lumen, which is a nutrient-rich environment with a competing microbiome (8), to severe nutritional restrictions and ROS attacks inside the so-called *Salmonella*-containing vacuole (SCV) during intracellular life within host cells (9, 10). Its versatile and robust metabolism (11) makes STM an ideal model organism to study the interconnection of metabolism and virulence functions.

To address the role of the TCA cycle in patho-metabolism of STM, we analyzed the effect of perturbations of the TCA cycle using a set of mutant strains each defective in one enzymatic step. Our former study indicated that TCA cycle perturbations induced in STM by oxidative stress result from damage of Fe-S cluster containing enzymes (12). Accordingly, a mutant strain deficient in all three fumarase isoforms (Δ*fumABC*) accumulated high amounts of TCA intermediate fumarate, but also showed the remarkable phenotype of increased phagocytosis by murine macrophages. These observations pointed towards a link between TCA cycle metabolite fumarate and celluar functions of STM.

The C_4_-dicarboxylate fumarate recently gained increasing interest due to various links between metabolisms and bacterial pathogenesis. In EHEC, fumarate is essential for full virulence in a *Caenorhabditis elegans* infection model where it regulates the expression of a tryptophanase by the transcription factor Cra (13). In *Mycobacterium tuberculosis*, fumarase deficiency was shown to be fatal due to protein and metabolite succination (14). Other studies demonstrated fumarate as factor that increases frequency of persister formation, or modulates motility and chemotaxis in *E. coli* (15–17).

In this work, we conducted metabolomics and proteomics to characterize the metabolic landscape of STM Δ*fumABC*. By this dual-omics approach, we elucidated a new example for the interplay between metabolism, and cellular functions and virulence in STM.

## Results

### Effects of TCA cycle enzyme deletion on the carbon metabolism of *S*. Typhimurium

For a global analysis of the effects of perturbations of the TCA cycle on patho-metabolism of STM, we generated a set of isogenic STM mutant strains, each defective in one reaction of the TCA cycle. Using this set of strains in comparison to STM WT, we performed metabolomics analyses of stationary cultures, grown 18.5 h in rich media (LB broth) and analysed samples as described before (12). Metabolomics revealed that the Δ*fumABC* strain, deficient in all fumarase isoforms, had a highly aberrant metabolic profile distinct from that of other mutant strains. Besides a strong accumulation of fumarate (115-fold compared to WT), STM Δ*fumABC* contained significantly increased amounts of glycolysis and PPP intermediates.

Moreover, the Δ*fumABC* strain exhibited increased levels of glucose-6-phosphate (G6P), fructose-6-phosphate (F6P) and sedoheptulose-7-phosphate (S7P), whereas all other mutant strains exhibited decreased or unchanged levels compared to WT (**Fig. 1**, **Table S 1**). This observation indicates distinct and unique impacts of the fumarase deletions on carbon flux. Only a mutant strain deficient in succinate dehydrogenase also showed a larger level of F6P, but not at the same extend as observed for Δf*umABC*. Furthermore, there was a strong accumulation of aspartate, likely arising from the large pool of fumarate by the action of aspartate ammonia-lyase AspA (**Table S 2**).

**Fig. 1.**
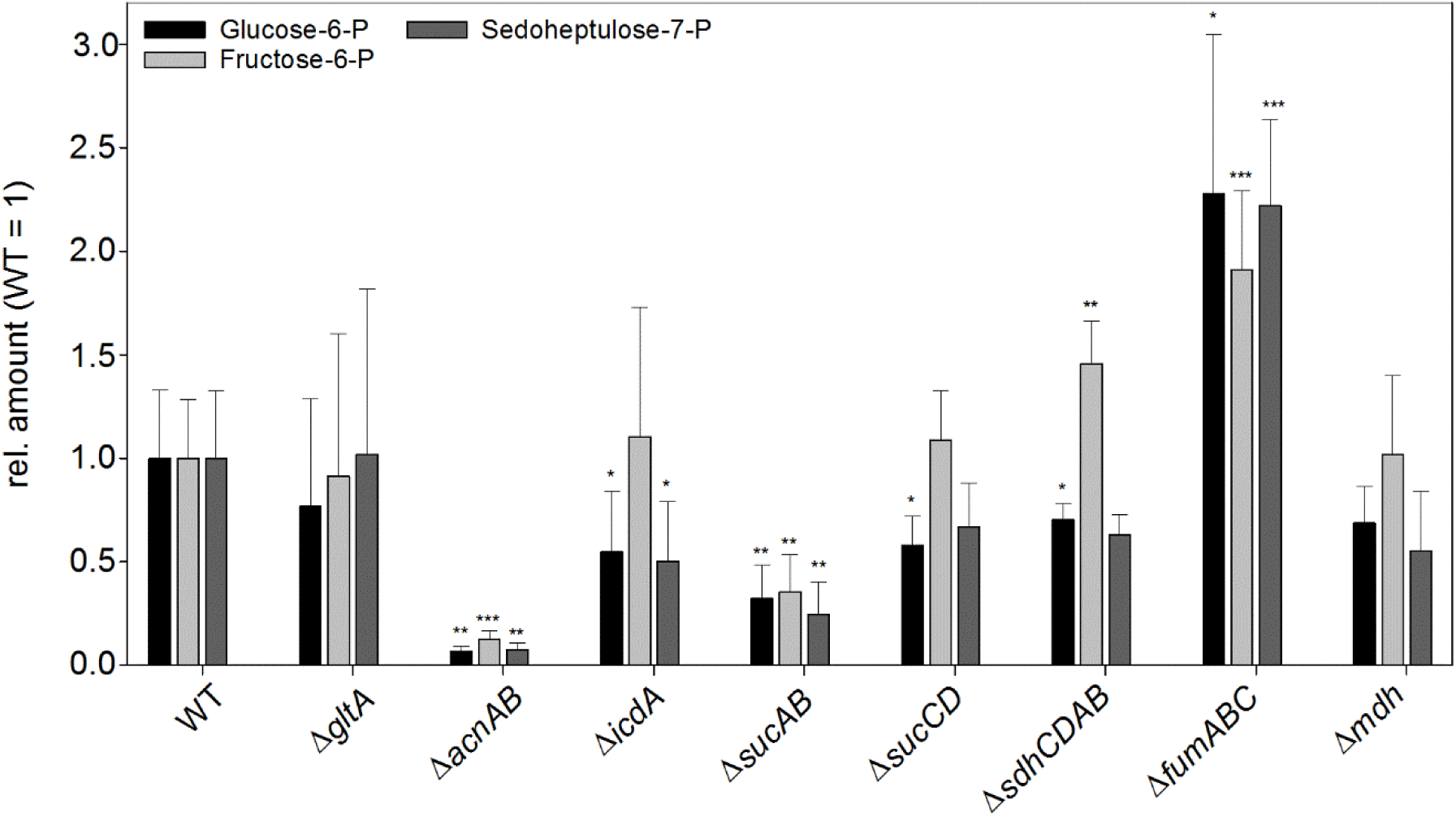
Defects in TCA cycle enzymes affect metabolite concentrations. STM WT and mutant strains defective in TCA cycle enzymes were grown aerobically in LB broth for 18.5 h at 37 °C. Cells were harvested, disrupted and metabolites extracted for subsequent GC-MS analyses. Metabolite concentrations were normalized to levels of WT, and means and standard deviations of at least four biological replicates are shown. Statistical analyses were performed by Student’s *t*-test and significances are indicated as follows: *, *p* < 0.05; **, *p* < 0.01; ***, *p* < 0.001.

In our previous analyses of ROS-induced damages of TCA cycle enzymes on STM patho-metabolism, we found that a mutant strain unable to detoxify endogeneously generated ROS was attenuated in intracellular proliferation. Surprisingly, this mutant strain was internalized by macrophages at higher rates than STM WT (12). Endogenous ROS cause damage of Fe-S cluster-containing TCA cycle enzymes, and also a Δ*fumABC* strain was internalized by macrophages at 15-fold higher rate compared to WT STM, without defects in intracellular proliferation. These observations point towards a link between the function of the TCA cycle and virulence properties of STM, which prompted us to characterize the STM Δ*fumABC* strain in detail.

### Quantitative proteomics and metabolic profiling reveal alterations in the central carbon metabolism of STM Δ*fumABC*

We first performed proteomic and metabolic profiling of STM WT and Δ*fumABC* strains after culture in rich media (LB broth) for 18.5 h and analyzed samples as described (12). As anticipated from genotype and fumarate accumulation, fumarases were not detected in the fumarase-deficient strain. We did not detetct changes in other TCA cycle intermediates (**Fig. 2**). However, we observed increased amounts of citrate synthase (GltA), aconitase B (AcnB), isocitrate dehydrogenase and α-ketoglutarate dehydrogenase component (SucA) by 2.05- to 2.72-fold. With respect to catabolism of hexoses, and in line with higher concentrations of G6P (2.28-fold) and slight increment of F6P (1.91-fold), increased amounts of the corresponding enzymes were detected in the Δ*fumABC* strain. Glucokinase (Glk), glucose-6-phosphate-isomerase (Pgi), phosphofructokinase A (PfkA) and phosphoglycerate mutase (GpmB) were only identified in Δ*fumABC*, and we determined 2.27- to 5.42-fold increased amounts of fructose-1,6-bisphosphatase class 1 (Fbp), fructose-bisphosphate-aldolase B (FbaB), glyceraldehyde-3-phosphate-dehydrogenase (GapA) and pyruvate kinase I (PykF). The increased amount of S7P can be correlated with higher amounts of glucose-6-phosphate-dehydrogenase (Zwf), ribulose-phosphate-3-epimerase (Rpe) and transketolase B (TktB), detected only in the proteome of Δ*fumABC*. Furthermore, transaldolase A (TalA) was increased 3.29-fold.

**Fig. 2.**
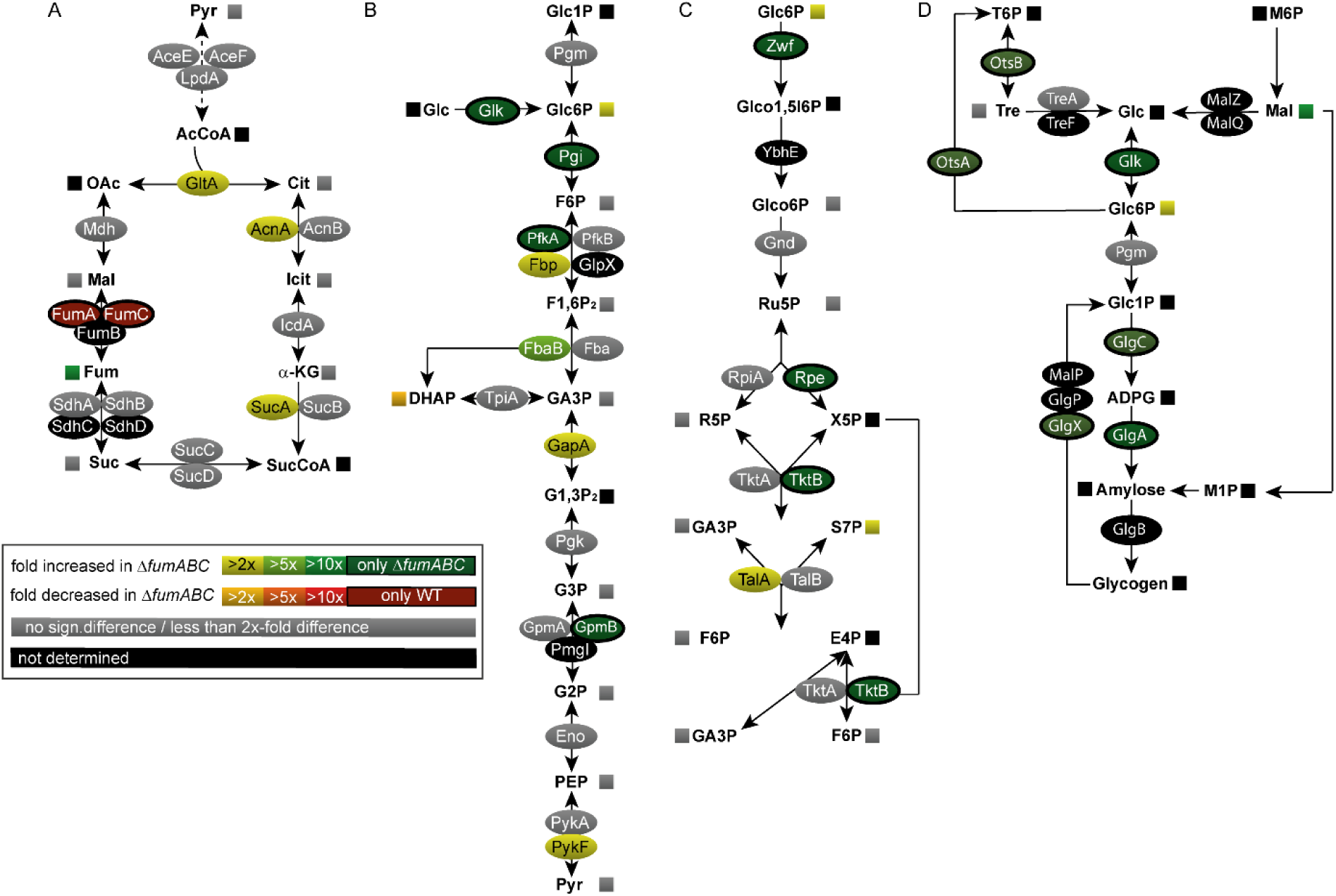
Deletion of fumarases leads to changes in carbon fluxes and amounts of metabolic enzymes. STM WT and Δ*fumABC* strains were grown aerobically in LB broth for 18.5 h at 37 °C. For the proteomic approach, harvested cells were lysed and proteins precipitated with 10% TCA. After trypsin digest, the peptides were analyzed by quantitative LC-MS^E^. For metabolomics analyses, harvested cells were disrupted and metabolites were extracted for GC-MS analysis. Heat map colors of oval symbols indicate relative changes in amounts of enzymes detected for Δ*fumABC* compared to WT. Heat map colors of square symbols indicate relative changes in amounts of metabolites determined in Δ*fumABC* compared to WT. Grey symbols indicate less than 2-fold or not significant differences in enzyme or metabolite amounts. Quantitative data are shown for TCA cycle (A), glycolysis (B), pentose phosphate pathway (C), and glycogen synthesis (D). Data represent means of at least four or three biological replicates for the metabolomics or proteomics analyses, respectively. Statistical analyses were performed by Student’s *t*-test and all data shown have significance differences between the two strains of *p* < 0.05 or lower.

In addition, we observed only in the proteome of STM Δ*fumABC* key enzymes of glycogen biosynthesis, i.e. glycogen synthase (GlgA), glucose-1-phosphate adenylyltransferase (GlgC), glycogen debranching enzyme (GlgX), trehalose-phosphate-synthase (OtsA), as well as trehalose-phosphate-phosphatase (OtsB) (**Fig. 2D**). Together with the detected accumulation of maltose (10-fold) and trehalose (2-fold), these data suggest an increased glycogen accumulation in STM Δ*fumABC* compared to the WT.

To test for increased glycogen storage, bacterial cultures grown on LB agar were treated with potassium iodine for glycogen staining (18). While STM WT was only lightly stained, the intense brown color of STM Δ*fumABC* colonies indicated high accumulation of glycogen (**Fig. 3C**). We next applied transmission electron microscopy (TEM) of ultrathin sections of STM WT (**Fig. 3A**) and Δ*fumABC* cells (**Fig. 3B**). Granular aggregates of low electron density were observed in the polar regions of STM Δ*fumABC*, but to a far lesser exteny in WT cells. Accordingly, enzymatic quantification revealed 12-fold increased glycogen content in STM Δ*fumABC* compared to WT (**Fig. 3D**). Complementation of STM Δ*fumABC* by plasmids harboring *fumAC* or *fumB* genes restored WT levels of glycogen (**Fig. S 1A**). These data indicate that fumarate accumulation in STM Δ*fumABC* is a key factor for biasing the glycogen metabolism towards altered carbon fluxes and increased glycogen storage.

**Fig. 3.**
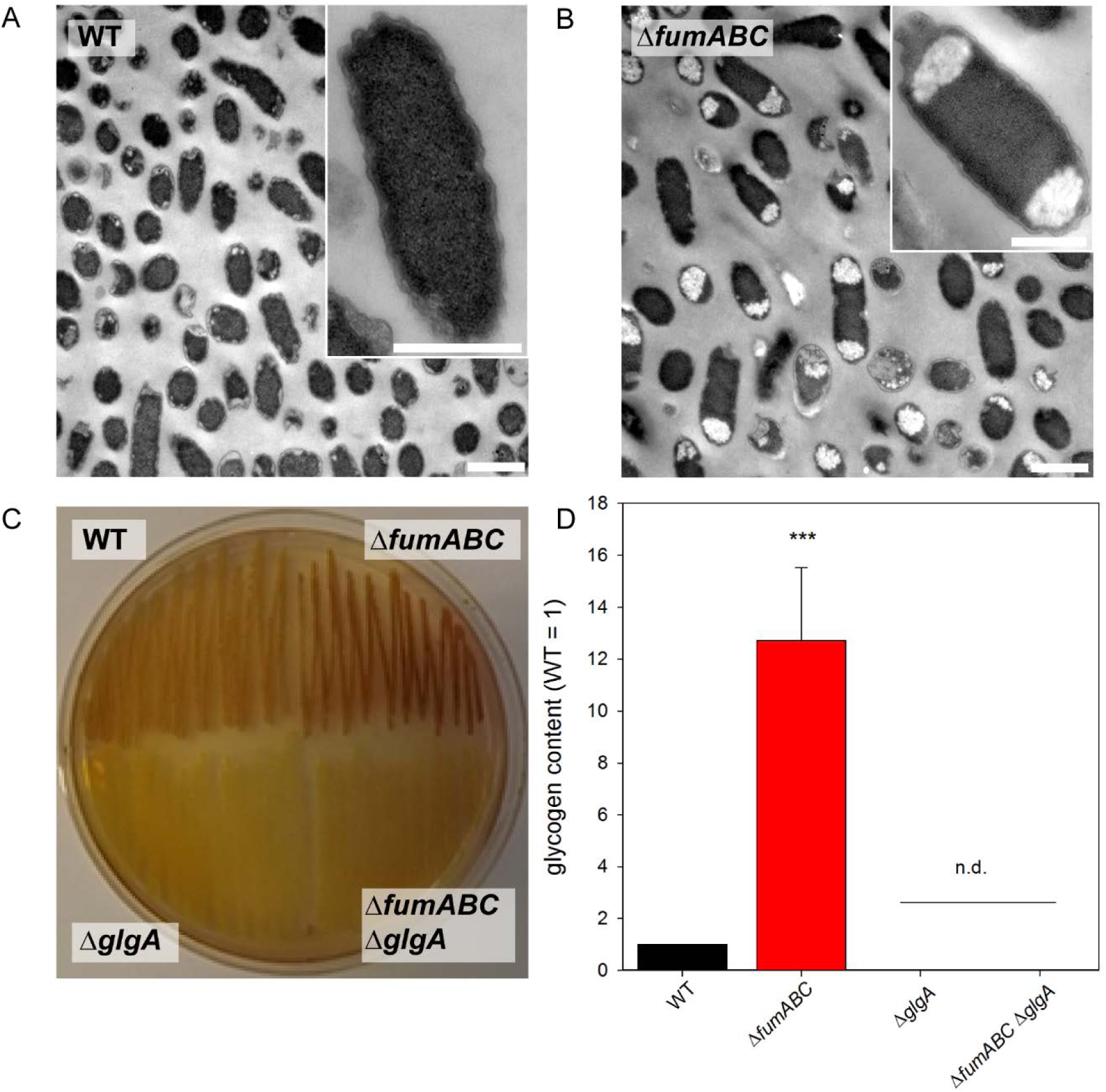
Deletion of fumarases leads to increased glycogen accumulation. STM WT (A) and Δ*fumABC* (B) strains were grown aerobically for 18.5 h at 37 °C in LB broth. Cells were fixed, dehydrated, resin embedded and ultrathin section were prepared for TEM. Massive accumulations of polymers in the polar regions of Δ*fumABC* cells were observed frequently. Scale bars, 1 µm (overview), 500 nm (detail). C) STM WT, Δ*glgA*, Δ*fumABC*, and Δ*fumABC* Δ*glgA strains* were grown on LB agar plates for 18.5 h at 37 °C. Potassium iodine staining was performed and brownish color indicates intercalation of iodine with glycogen. D) Quantification of glycogen contents of STM strains grown aerobically for 18.5 h in LB broth. Glycogen was degraded to glucose monomers using amyloglucosidase, and resulting glucose was phosphorylated to G6P. G6P was oxidized by G6P dehydrogenase in the presence of NAD, being reduced to NADH. Glucose concentrations were proportional to OD_340_. By subtraction of free glucose concentrations (sample without amyloglucosidase) from total glucose concentrations, glycogen amounts were quantified. Glycogen concentrations were normalized to WT (=1), error bars represent standard deviations of four biological replicates. n.d., not detected. Statistical analyses were performed by Student’s *t*-test and significances are indicated as follows: ***, *p* < 0.001.

### Deletion of glycogen synthase GlgA decreases amounts of G6P, F6P and S7P in *Salmonella* WT and Δ*fumABC* strains

To further investigate the connection of glycogen biosynthesis and fumarate accumulation, we blocked glycogen synthesis by deletion of *glgA*, which encodes the glycogen synthase, in the Δ*fumABC* mutant resulting in the STM Δ*fumABC* Δ*glgA* double mutant. We verified the loss of glycogen production in the *glgA*-deficient strain with potassium iodine staining (**Fig. 3C**) and TEM analyses (**Fig. S 2**) as before and were able to restore the original phenotype by complementation with a plasmid harboring *glgA* (**Fig. S 1B**).

Subsequently, we performed quantitative comparative proteomics and metabolomics of STM Δ*fumABC* Δ*glgA* and compared the obtained profiles with those of STM Δ*fumABC* (**Fig. 4**, **Table S 1, Table S 3**). Deletion of glycogen synthase did not affect amounts of metabolic enzymes in glycolysis, PPP and TCA cycle, but decreased the abundance of glucose-1-phosphate adenylyltransferase GlgC, an enzyme catalyzing the synthesis of ADPG. Metabolite analyses by GC-MS revealed strong decrease of G6P, F6P and S7P if *glgA* is deleted **Fig. 4 E**. Furthermore, the amount of trehalose was increased by 30%, while amounts of maltose were 100-fold reduced in STM Δ*fumABC* Δ*glgA*.

**Fig. 4.**
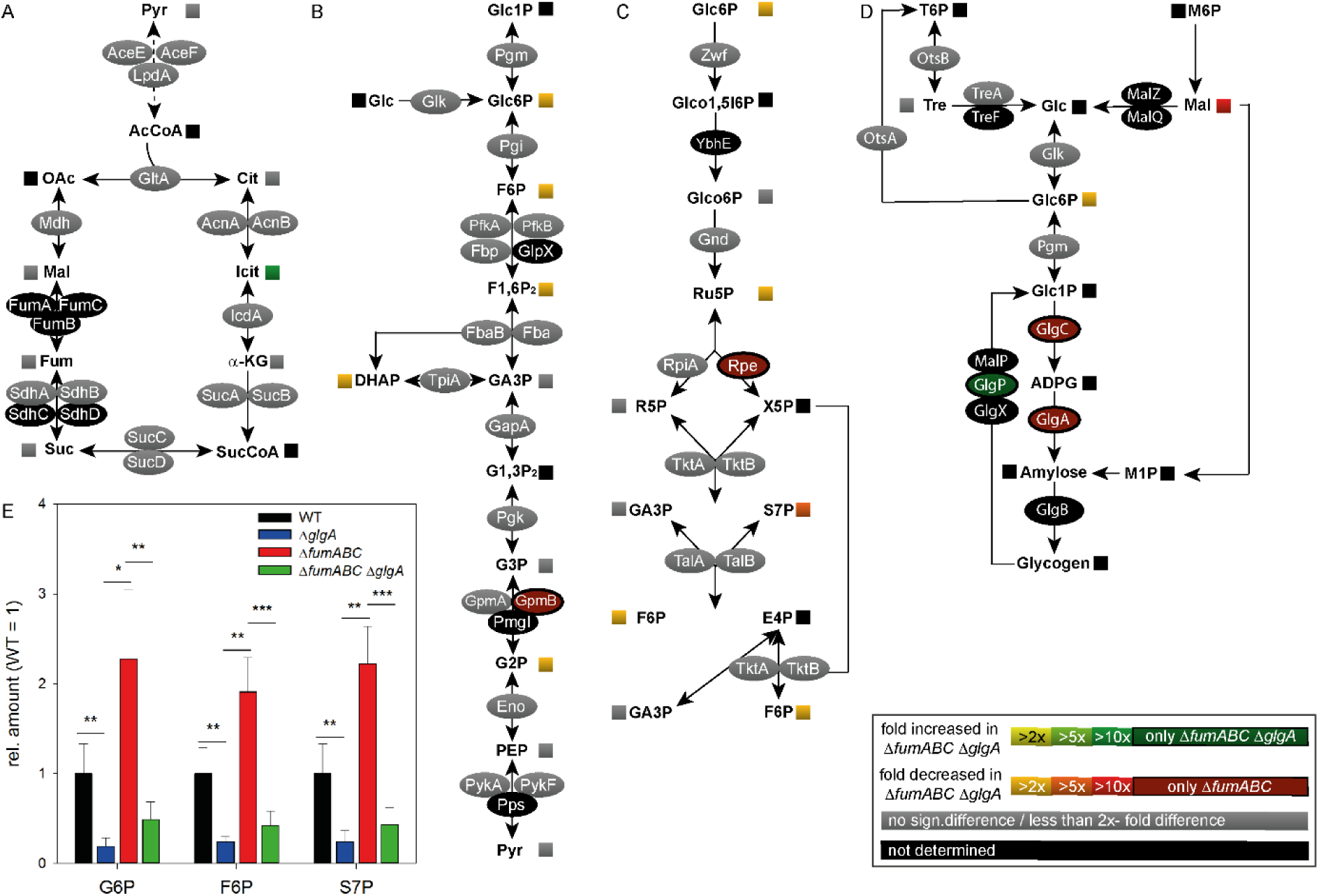
Deletion of glycogen synthase in a Δ*fumABC* strain restores carbon fluxes without changes in amounts of metabolic enzymes. STM Δ*fumABC* and Δ*fumABC* Δ*glgA* were grown aerobically in LB broth for 18.5 h at 37 °C. Analyses of metabolic enzymes and metabolites were performed as described for Fig. 2 and comparison of STM Δ*fumABC* Δ*glgA* to Δ*fumABC* are shown for TCA cycle (A), glycolysis (B), pentose phosphate pathway (C), and glycogen synthesis (D). Data represent means of at least four or three biological replicates for the metabolomics or proteomics analyses, respectively. The concentrations of metabolites glucose-6-phosphate (G6P), fructose-6-phosphate (F6P) and sedoheptulose-7-phosphate (S7P) were determined and normalized to WT (=1) (E). Statistical analyses were performed by Student’s *t*-test and all data shown have significance differences between the two strains of *p* < 0.05 or lower: *, *p* < 0.05; **, *p* < 0.01; ***, *p* < 0.001.

We conclude that altered fluxes through glycolysis and PPP in a fumarase-deficient strain are induced by increased glycogen synthesis. Abrogation of storage compound synthesis by *glgA* knockout normalized metabolite levels, due to modified enzyme activities and regulative mechanisms, rather than altered protein amounts.

### Fumarate-induced stringent response influences *Salmonella* physiology

The amount of stored glycogen is dependent on the abundance of synthesis enzymes (19), and glycogen synthesis in STM is mainly mediated by enzymes GlgA and GlgC (20). In *E. coli*, the main regulators for *glgA* and *glgC* transcription are the alarmones ppGpp and pppGpp (further referred to as (p)ppGpp) (21), which are induced during nutrient starvation by stringent response mediators RelA and SpoT. To elucidate whether STM Δ*fumABC* has an enhanced stringent response compared to STM WT, we made use of a dual-color reporter plasmid for relative quantification of *wraB* (= *wrbA* in *E. coli*) expression, which was recently used to determine the (p)ppGpp levels in *E. coli* (22). We introduced the P*_wraB_*::sfGFP reporter plasmid into STM WT, Δ*fumABC*, Δ*fumABC* Δ*glgA*, and as negative control into STM Δ*relA* Δ*spoT*, a mutant strain deficient in (p)ppGpp synthesis (23), and analyzed the expression by flow cytometry (**Fig. 5**). To test reporter performance, stationary LB broth cultures of STM WT were sub-cultured in defined PCN minimal media with or without supplemention by casamino acids (**Fig. 5A**). Indeed, WT grown without additional source of amino acids showed a higher sfGFP signal intensity compared to STM WT grown with amino acid supplementation, indicating higher (p)ppGpp levels.

**Fig. 5.**
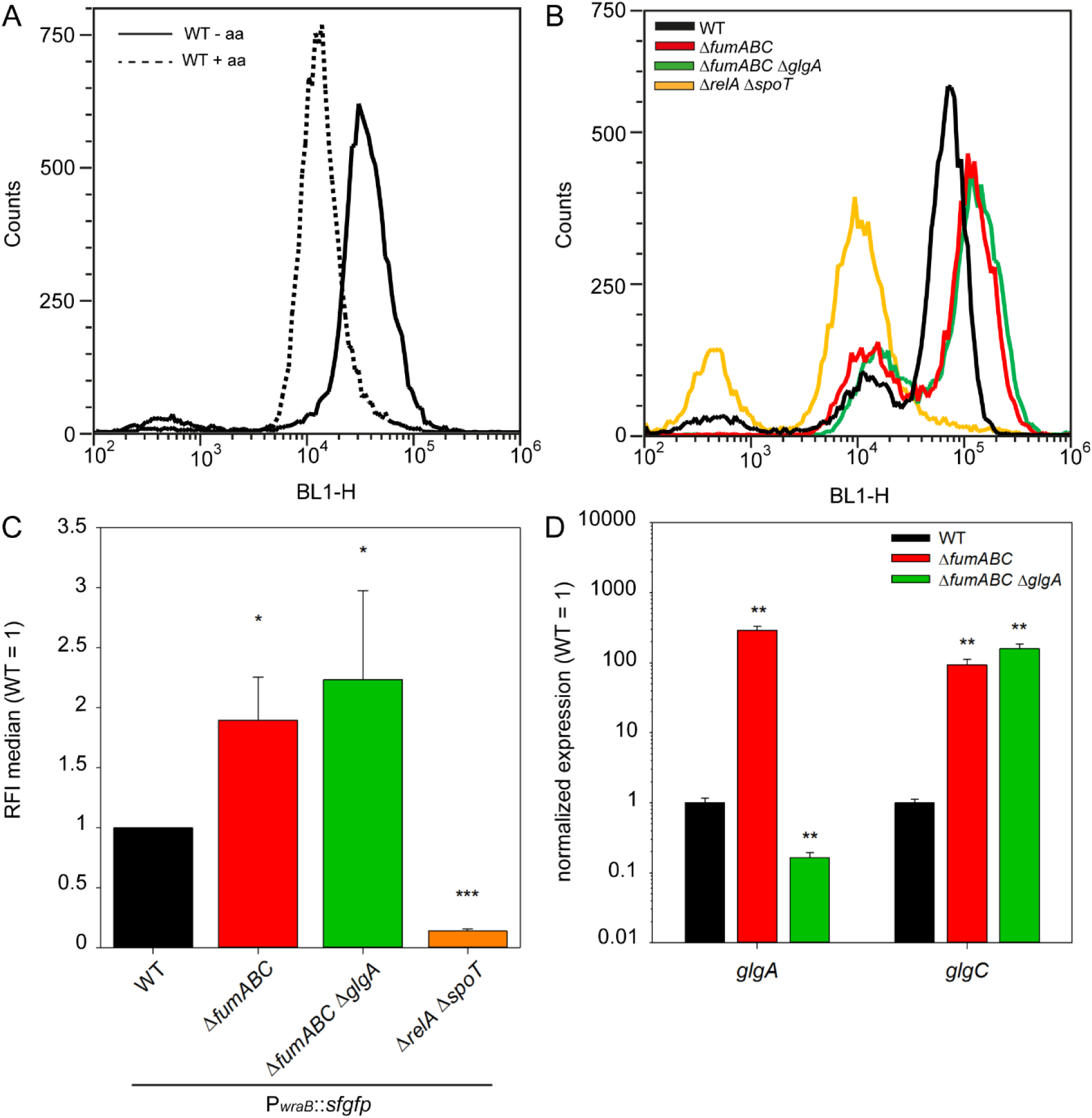
Differential expression of glycogen synthesizing enzymes due to increased (p)ppGpp levels in Δ*fumABC*. A) STM WT harboring a dual color fluorescence reporter for *wraB* was cultured in LB o/n and subcultured in minimal medium with or without amino acid (= aa) supplementation (dashed or undashed line, respectively). After 3 h of growth cells were subjected to flow cytometry and sfGFP fluorescence intensity (BL1-H) recorded. Shown is one representative of three independent biological replicates. B) Representative data of WT, Δ*fumABC*, Δ*fumABC* Δ*glgA* and Δ*relA* Δ*spoT* strains harboring the *wraB* reporter grown o/n in LB broth. C) Medians of relative sfGFP fluorescence intensities of strains mentioned in (B). Data were normalized to WT (=1) and represent average values and standard deviation of three biological replicates. D) WT, Δ*fumABC* and Δ*fumABC* Δ*glgA* strains were cultured o/n in LB broth, RNA was extracted and used for cDNA synthesis and consecutive qPCR experiments. 16s rRNA expression levels were used for normalization. Depicted are the expression levels normalized to WT (=1) of *glgA* and *glgC*. Shown is one representative assay of three independent biological replicates, consisting each of three technical replicates. Statistical analyses were performed by Student’s *t*-test and significances are indicated as follows: *, *p* < 0.05; **, *p* < 0.01; ***, *p* < 0.001.

Next, we determined sfGFP signal intensities of STM WT, Δ*fumABC* and Δ*fumABC* Δ*glgA* harboring the respective reporter plasmid cultured in LB broth for 18.5 h as described before. Quantification of sfGFP intensity revealed higher values for STM Δ*fumABC* and Δ*fumABC* Δ*glgA* compared to STM WT, whereas the negative control STM Δ*relA* Δ*spoT* exhibited the lowest signal intensities (**Fig. 5BC**). Additionally, transcript levels of *glgA* and *glgC* were determined (**Fig. 5D**). Strongly enhanced expression of *glgA* and *glgC* was detected for Δ*fumABC* compared to WT. For STM Δ*fumABC* Δ*glgA*, we only detected background signals for *glgA*, but still highly increased expression levels of *glgC* compared to WT. In addition, glycogen accumulation in STM Δ*fumABC* was eliminated by further deletion of *relA* and *spoT* (**Fig. S 3**). Thus, we propose that Δ*fumABC* enforces glycogen synthesis as consequence of an early and strong stringent response, leading to high (p)ppGpp levels, which in turn raises the transcript and protein levels of GlgA and GlgC.

### Altered amounts of chemotaxis proteins in fumarase–deficient STM lead to increased CCW flagella rotation

Accumulation of (p)ppGpp can negatively affect motility, as recently described for *E. coli* (24). To explore this potential link, we analyzed proteomic data for modulation of chemotaxis and motility-related proteins (**Fig. 6A**). Decreased amounts of methyl-accepting chemotaxis proteins (MCP) and increased abundance of CheY, CheZ and CheW (2.14-3.86-fold) were detected in STM Δ*fumABC* compared to WT. In addition, CheB was only found in STM Δ*fumABC*. For STM Δ*fumABC* Δ*glgA*, a restoration of chemotaxis protein levels was detected for CheY. However, CheY abundance was still lower compared to STM WT (**Fig. 6B**).

**Fig. 6.**
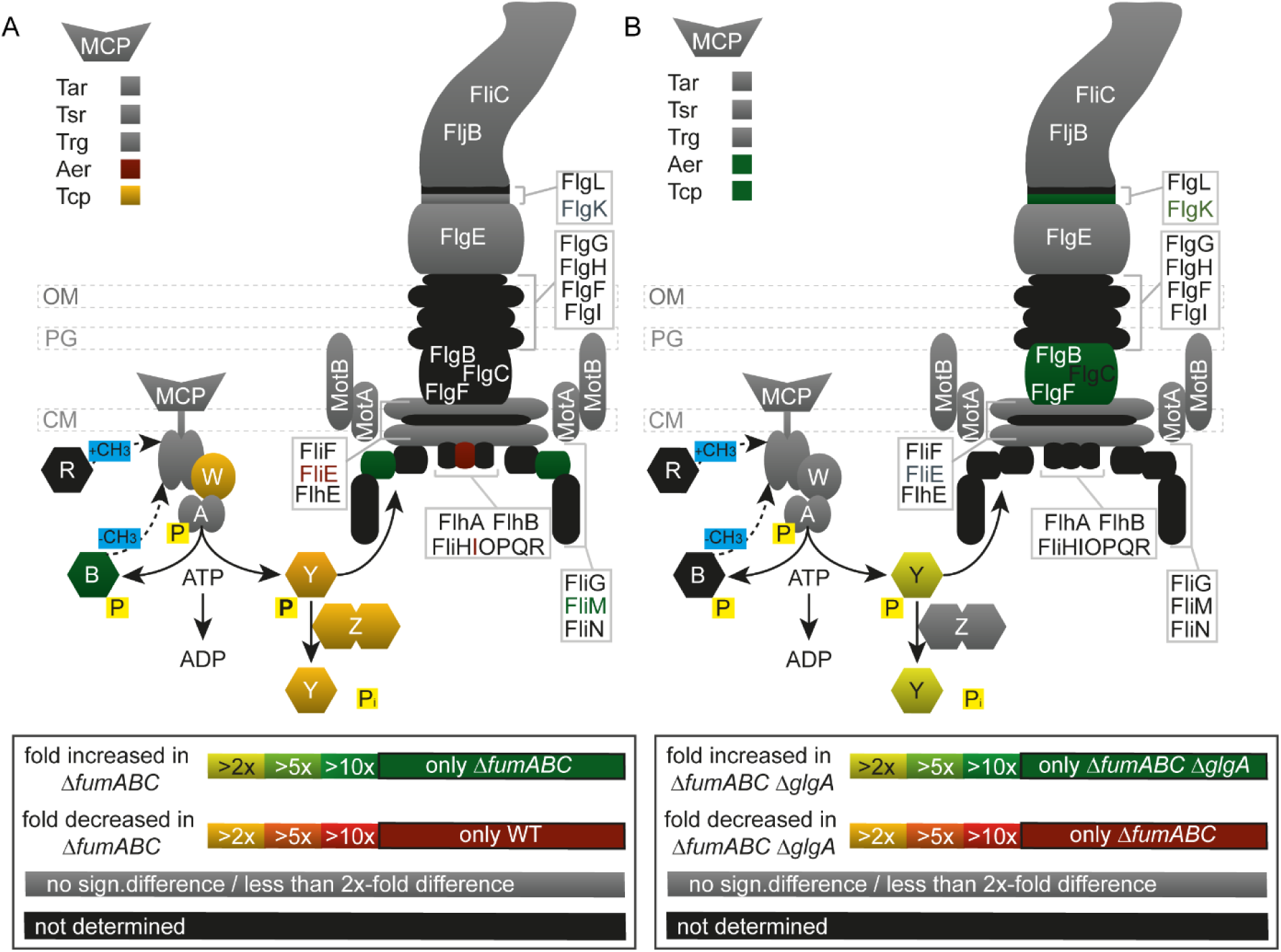
Deletion of fumarases affects amounts of flagellar and chemotaxis proteins, which is partly restored by deletion of *glgA*. STM WT, Δ*fumABC and* Δ*fumABC ΔglgA* strains were grown aerobically in LB broth for 18.5 h at 37 °C. For proteomic analyses, cells were harvested, lysed, and proteins precipitated by 10% TCA. After trypsin digestion, peptides were analyzed by quantitative LC-MS^E^. Relative changes in protein abundance detected for (A) Δ*fumABC* compared to WT, and (B) Δ*fumABC ΔglgA* to Δ*fumABC* are indicated by color heat maps. Symbols in the upper left indicate abundance of methyl-accepting chemotaxis proteins (MCP). Symbols in the lower left corner represent chemotaxis proteins, letters indicate subunits (e.g. Y for CheY). P indicates phosphorylation. Data represent means of at least three biological replicates. Statistical analyses were performed by Student’s *t*-test and all shown data illustrate statistical significant differences between the two strains (*p* < 0.05).

The amount of CheY influences the number of switching events of flagella rotation direction (25). Thus, STM Δ*fumABC* might show an altered swimming behavior and we analyzed swim patterns of bacteria grown over night in rich medium (**Fig. 7A**). Counterclockwise (CCW) flagella rotation bundles flagella and results in straight swimming, while clockwise (CW) rotation leads to tumbling (26). STM WT showed short swimming paths alternating with tumbling, whereas STM Δ*fumABC* exhibited highly prolonged swimming paths and reduced tumbling events. Furthermore, the number of motile bacteria was higher compared to WT. The motility patterns of Δ*fumABC* and Δ*fumABC* Δ*glgA* were similar.

**Fig. 7.**
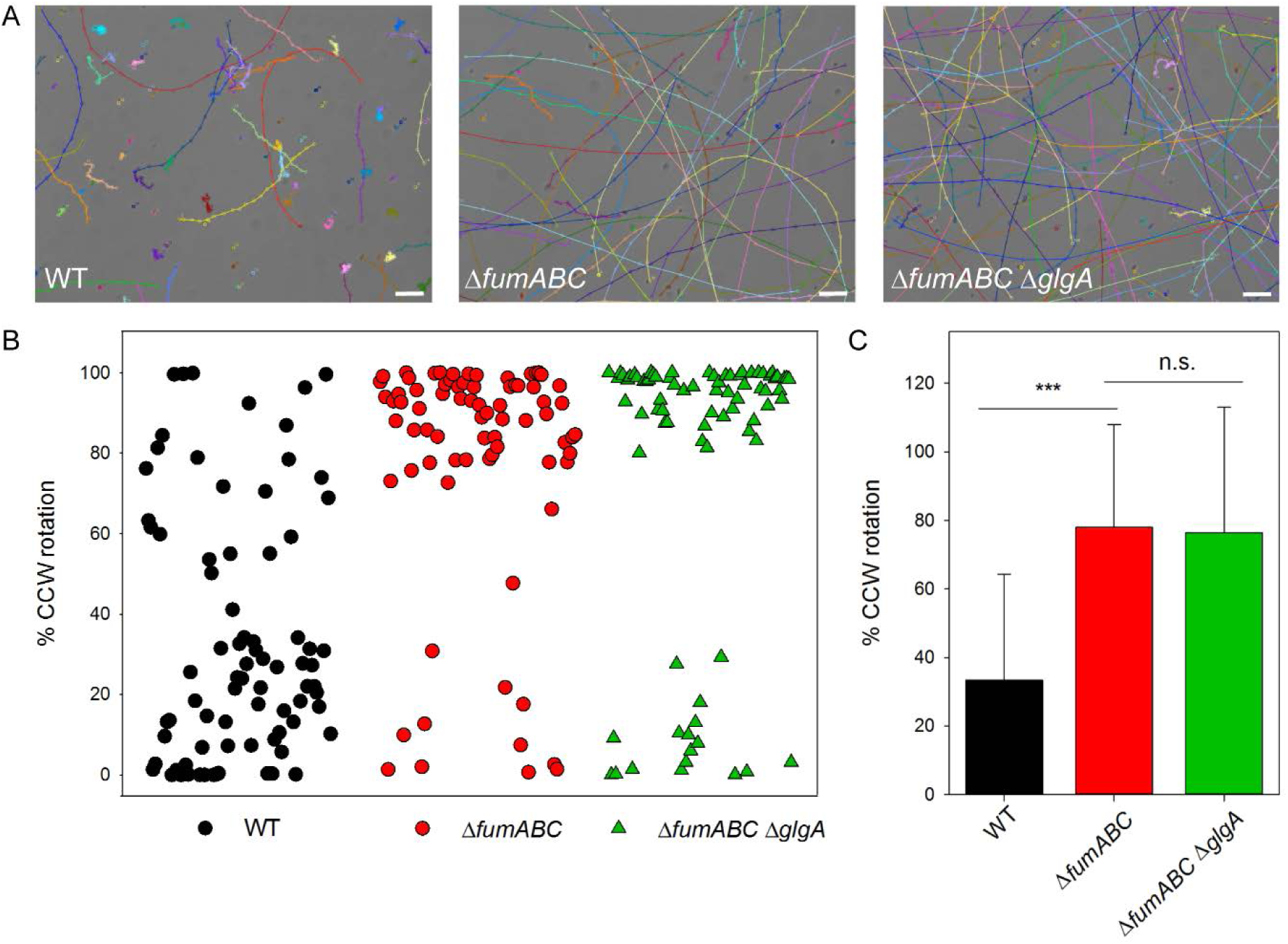
Fumarase deletions increase CCW flagella rotation and running motility in a *glgA*-independent manner. A) STM WT and mutant strains were cultured in LB broth for 18.5 h at 37 °C and diluted 1:20 in PBS. Aliquots of the cultures were spotted onto microscope slides and bacterial motion was recorded at 14 frames/s for 100 frames. For manual path tracking, the ImageJ plugin MTrackJ was used. Paths of individual STM WT, Δ*fumABC* or Δ*fumABC* Δ*glgA* cells are indicated by various colors. Scale bars, 10 µm. B) Bacteria were cultured for 18.5 h in LB, diluted 1:100 in PBS, subjected to shear force to remove flagellar filaments, and bound to polystyrene-coated coverslips. Rotating cells were selected and rotation direction was recorded using the Axio Observer microscope with an AxioCam CCD camera (Zeiss) for periods of 18 s. Rotation analyses were performed using the tool SpinningBug Tracker. By detection of the angle between the rotating bacteria and a reference axis, the rotation direction was calculated. Each dot represents the analysis of one bacterial cell. The experimental setup and definitions are illustrated in **Fig. S 5**. C) Quantification of CCW bias of single STM cells. Means and standard deviations are shown for at least 75 cells per strain. Statistical analyses were performed by Rank Sum Test and significances are indicated as follows: ***, *p* < 0.001; n.s., not significant.

To further analyze flagella switching from CCW to CW rotation, we performed flagella rotation analyses of STM WT, Δ*fumABC* and Δ*fumABC* Δ*glgA* grown in rich medium by microscopic inspection of single bacterial cells fixed by one flagellum to a polystyrene-coated coverslip (27) (**Fig. 7BC**). We observed a statistically significant increase of CCW flagella rotation for STM Δ*fumABC*. Whereas STM WT had an average proportion of CCW rotation of 33%, the Δ*fumABC* strain spent 78% of time in CCW flagella rotation. Although STM Δ*fumABC* Δ*glgA* exhibited partly normalized amounts of the chemotaxis protein CheY, there was still an increased proportion of CCW flagella rotation comparable to that of STM Δ*fumABC*. Furthermore, the swimming behavior was not altered by *glgA* deletion, indicating that the amount of CheY necessary for normalization of switching events was not achieved in STM Δ*fumABC* Δ*glgA*.

Thus, we conclude that fumarase deletion in STM leads to a down-regulation of chemotaxis proteins and by this to enhanced CCW flagella rotation.

### The increased phagocytic uptake of fumarase-deficient STM is due to enhanced CCW flagella rotation and partially depends on glycogen synthesis

Since bacterial motility can increase uptake of pathogenic bacteria by host cells (28–31), we hypothesized that the observed enhanced uptake of *fumABC* mutant strains by RAW264.7 macrophages (12) could be caused by increased CCW flagella rotation. To test this hypothesis, we introduced additional delection of chemotaxis genes *cheY* or *cheZ* in the mutant strain. Whereas *cheY* deletion strains are locked in CCW flagella rotation, Δ*cheZ* mutant strains are mainly locked in the CW state (32). The combination of Δ*cheY* and Δ*fumABC* did not alter phagocytic uptake, while the combination of Δ*cheZ* and Δ*fumABC* showed uptake only 3.15-fold higher than WT (**Fig. 8**).

**Fig. 8.**
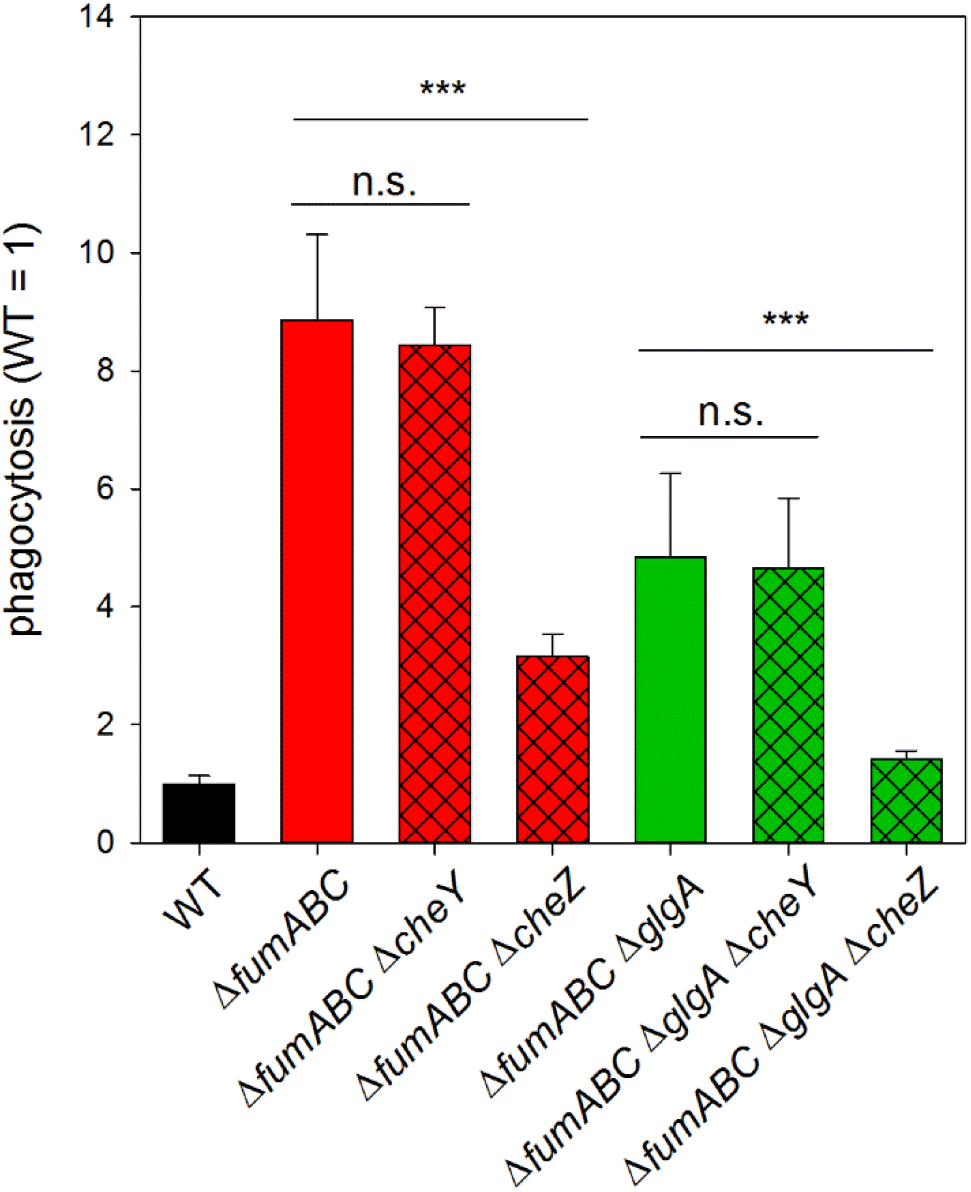
Increased phagocytic uptake of fumarase deletion strains is dependent on CCW flagella rotation and glycogen synthesis. STM WT and various mutant strains as indicated were grown for 18.5 h in LB broth and used to infect RAW264.7 macrophages at an MOI of 1. Infection was synchronized by centrifugation for 5 min., followed by incubation for 25 min. at 37 °C. Next, non-internalized bacteria were removed by washing and treatment with gentamicin at 100 µg/ml for 1 h and 10 µg/ml for the remaining time. Cells were lysed 2 h after infection by addition of 0.1% Triton X-100 and lysates were plated onto MH agar plates to determine the CFU of internalized bacteria. Phagocytosis rates were determined as percentage of internalized bacteria in dependence of the used inoculum. Values were normalized to WT (=1), and means and standard deviations of three technical replicates are shown. Statistical analyses were performed by Student’s *t*-test and significances are indicated as follows: ***, *p* < 0.001; n.s., not significant.

Deletion of glycogen synthase partially normalized CheY levels, but not the duration of CCW flagella rotation in STM Δ*fumABC*. Thus, we expected an increased phagocytic uptake of the Δ*fumABC* Δ*glgA* double mutant as well. However, phagocytosis of STM Δ*fumABC* Δ*glgA* was 6.9-fold increased compared to WT, but significantly lower than uptake of STM Δ*fumABC* (**Fig. 8B**). Complementation by plasmid-borne *glgA* again increased levels of phagocytosis (**Fig. S 4**). The *cheY* deletion did not change phagocytic uptake of STM Δ*fumABC* Δg*lgA*, while phagocytosis of STM Δ*fumABC* Δ*glgA* Δ*cheZ* was reduced (**Fig. 8**). These results demonstrate that high phagocytosis of fumarase deletion strains is due to CCW bias of flagella rotation and is partially dependent on glycogen synthesis.

In order to elucidate which factors reduce the phagocytic uptake of STM Δ*fumABC* Δ*glgA* compared to Δ*fumABC*, we analyzed further characteristics of swimming behavior of both mutant strains. The frequency of switching events within 1,000 frames (17.71 s) was determined and a switching event occurred if the flagella rotation direction changed from CW to CCW or *vice versa* (**Fig. S 5**). Compared to WT (median = 31 events), the switching rate was reduced in STM Δ*fumABC* (median = 20 events), but not in a significant manner (**Fig. 9A**). Even stronger reduction of switching events was determined for STM Δ*fumABC* Δ*glgA* (median = 10 events). Additionally the number of pauses, defined as rotation of the bacterial body of less than 5°/frame, was analyzed (**Fig. 9B**). Comparable to the number of switching events, WT had the highest number of pauses (median = 170.5), followed by STM Δ*fumABC* (151.5), but again there is no statistical significant difference between these two strains. A stronger reduction was observed for STM Δ*fumABC* Δ*glgA*; here the number of pauses within 1,000 frames was reduced to 89.

**Fig. 9.**
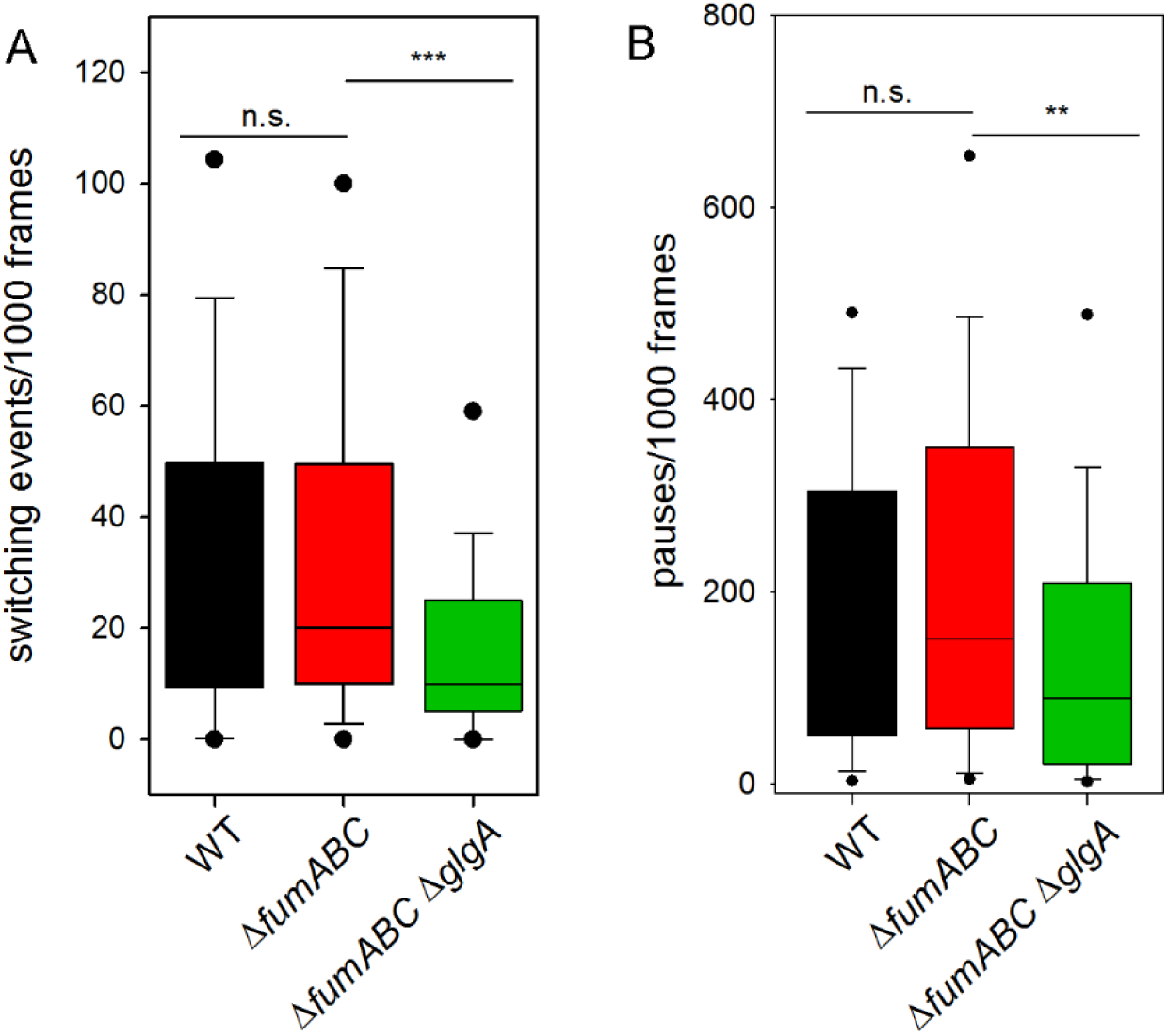
Deletion of fumarases and glycogen synthase effect the number of flagellar switching events and pauses. A) Distribution of the number of switching events within 1,000 frames (total of 17.54 s) was calculated using flagella rotation analysis (Fig. 7). B) Distribution of pauses within 1000 frames. Pause is defined as movement of the bacterial body less than 5°/frame. Values within the 5/95% percentiles are excluded. Calculations are based on at least 75 analyzed bacteria. Statistical analyses were performed by Rank Sum Test and significances are indicated as follows: **, *p* < 0.01;***, *p* < 0.001; n.s., not significant.

In concusion, STM Δ*fumABC* showed strongly increased CCW bias and less switching events than STM WT. These factors influence the interaction with host cells, such as increasing phagocytic uptake by macrophages. Further deletion of *glgA* in the Δ*fumABC* strain did not reduce time spent in CCW flagella rotation, but decreased the number of switching events, resulting in reduced phagocytic uptake.

## Discussion

Our work investigated the effect of perturbation of the TCA cycle of STM on basic cellular functions and patho-metabolism. By deploying proteomics and metabolomics, we determined that defects in fumarases biased carbon fluxes towards enhanced glycogen synthesis, likely due to elevated (p)ppGpp levels in the mutant strain. Furthermore, proteomics revealed reduced abundances of chemotaxis proteins in STM Δ*fumABC*. Analysis of flagella rotation and swim patterns showed increased CCW bias, raising the contact frequency of STM and host cells, thus leading to enhanced phagocytic uptake by macrophages. Deletion of glycogen synthase GlgA relieved the metabolic perturbations, but not the aberrant motility phenotype. However, phagocytic uptake was decreased. These findings are in line with a previous study that analyzed the impact of deletions of TCA cycle enzymes harboring Fe-S clusters on patho-metabolism of STM (12). For Δ*fumABC*, we measured increased abundance of glycolysis and PPP metabolites, and an elevated phagocytic uptake by RAW264.7 macrophages.

Our metabolomics data demonstrated higher accumulation of G6P, F6P and S7P for STM Δ*fumABC* compared to WT, and deletion of glycogen synthase again normalized the metabolic flux through glycolysis and PPP (**Fig. 4**). Thus, the increased concentrations of these metabolites were caused by enhanced glycogen synthesis in STM Δ*fumABC* due to changes in carbon fluxes. Accumulation of the respective metabolites was also observed for *E. coli* with truncated CsrA, the main component of the carbon storage system (33, 34). As *csrA* deletion strains accumulate high amounts of glycogen as well, our results indicate that the observations obtained for *E. coli* Δ*csrA* are also consequence of the massive remodeling of the carbon metabolism due to enhanced glycogen synthesis. However, a role of CsrA was not only reported in context of post-transcriptional regulation of carbon metabolism, and in particular glycogen metabolism, but also for chemotaxis proteins, flagella subunits and proteins involved in virulence functions (35, 36). Thus, the involvement of CsrA as inducer of phenotypes of STM Δ*fumABC* is conceivable. While glycogen accumulation indicates very low levels of CsrA, mutant strains with truncated CsrA showed increased levels of Pgm and reduced levels of especially PfkA in *E. coli* (33), observations which are contradictory to our results. However, most studies on *csrA* mutant strains were performed with bacteria grown in minimal media, or at early growth phases (33). Thus, we cannot exclude a role of CsrA in the enhancement of glycogen synthesis for STM Δ*fumABC*, yet we do not expect CsrA to be sole regulating factor. In contrast, (p)ppGpp was shown to be the most important factor influencing glycogen synthesis, at least in *E. coli* (37). (p)ppGpp is known to enhance the expression of *glgA* and *glgC*, but not *glgB* during stringent response (19). Indeed, we detected GlgA and GlgC only in STM Δ*fumABC* (**Fig. 2D**). Using a dual-color reporter system with P*_wraB_* controling sfGFP expression, we detected increased fluorescence intensities for STM Δ*fumABC* and Δ*fumABC* Δ*glgA* compared to WT. The promoter of *wrbA* was used in several studies for the indirect quantification of (p)ppGpp (22, 38). Furthermore, by proteomic analyses we detected increased abundances of WrbA in Δ*fumABC* (3.78-fold, see Table S2), supporting our results obtained by flow cytometry. Taken together, we hypothesize that a fumarase deletion strain increases *glgA* and *glgC* expression in a (p)ppGpp-dependent manner.

The main inducing factors for (p)ppGpp synthesis by RelA and SpoT are amino acid and carbon source limitations (39). Using LB broth, amino acid limitations are unlikely at early growth phase. Several studies showed that increase of (p)ppGpp levels can be induced by diauxic shifts, for example from glucose to succinate (40). Considering the high accumulation of fumarate, the use of the TCA cycle intermediate as carbon source is conceivable. An indicator for this model is the slightly increased abundance of aspartase AspA in STM Δ*fumABC* (1.5-fold), catalyzing the reversible reaction from fumarate and ammonia to aspartate (41). Indeed, metabolomic data showed a 10-fold higher amount of aspartate in the mutant strain, which serves as substrate for a range of metabolic pathways (42). Furthermore, two studies indicated that high fumarate accumulation led to use of fumarate as alternative electron acceptor, despite presence of oxygen (15, 43). However, our proteomic data gave no hints for fumarate respiration (i.e. fumarate reductase FrdABCD) in the mutant strain, but rather indicated utilization of fumarate as carbon source. Fumarate metabolism possibly leads to a physiological situation similar to exponential to stationary phase transition, and therefore increased (p)ppGpp levels, as discussed for *E. coli* (15, 22).

Absence of fumarases led to enhanced CCW flagella rotation, prolonged phase of running movement, resulting in increased uptake by RAW264.7 macrophages (**Fig. 8**). The impact of CCW flagella rotation during the infection process was discussed in several prior publications (28–30). In these studies, CCW flagella rotation and the resulting smooth swimming phenotype were linked to enhanced frequencies of bacterial contact with host cells, prolonged duration of adhesion, and increased numbers of phagocytic uptake events. Further deletion of *glgA* in STM Δ*fumABC* partly restored CheY levels and we observed reduced uptake of STM Δ*fumABC* Δ*glgA* by macrophages. As we determined a strongly decreased number of switching events for the *glgA*-deficient strain, but high frequency of phases of CCW flagella rotation, the logical consequence is that duration of phases of CW flagella rotation after switching events are longer for STM Δ*fumABC* Δ*glgA* than for Δ*fumABC*. This effect might be accompanied by the reduced number of pause events observed for STM Δ*fumABC* Δ*glgA* in comparison to Δ*fumABC* and WT and could lead to changes in frequency or duration of contacts between STM and host cells. To conclude, our results demonstrate that accumulation of fumarate due to fumarase deletion leads to induction of glycogen synthesis by enhanced (p)ppGpp concentrations (**Fig. 10**). This might be triggered by utilization of fumarate as carbon source, causing a exponential to stationary phase transition-like physiological state during early stationary growth phase. Additionally, we revealed that the increased phagocytic uptake of the fumarase deletion strain is caused by enhanced CCW flagella rotation, which is the consequence of reduced CheY abundance. Further deletion of *glgA* normalized metabolic fluxes and restored abundance of the chemotaxis protein in part, but did not change CCW bias of flagella rotation. However, *glgA* deletion led to reduced phagocytic uptake by RAW264.7 macrophages, possibly due to prolonged periods of CW flagella rotation. Our work demonstrates that perturbations of the carbon fluxes in the TCA cycle lead to dramatic changes in STM physiology and affect the interaction of this pathogen with host cells.

**Fig. 10.**
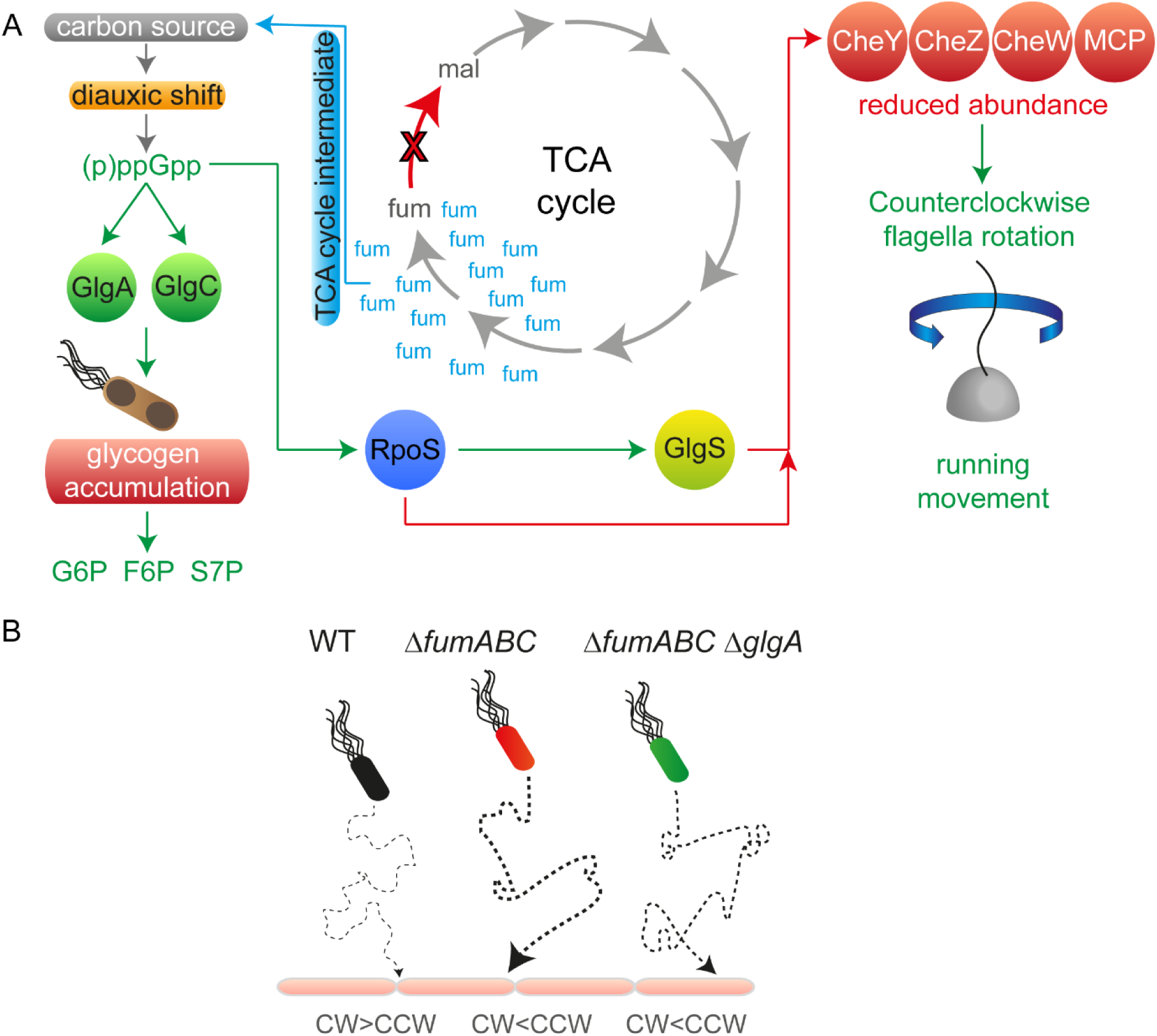
Model summarizing effects of fumarase deletion on STM patho-physiology. A) Fumarase deletion leads to fumarate accumulation. If present in high concentrations it is used as carbon source, which induces a kind of diauxic shift and increased (p)ppGpp concentrations. The alarmone accelerates the expression of glycogen synthesizing enzymes and glycogen accumulation occurs. This metabolic shift is accompanied by increased concentrations of glycolytic and PPP intermediates. (p)ppGpp strengthens RpoS biosynthesis, which in turn enhances GlgS abundance. GlgS and RpoS reduce expression of chemotaxis genes, leading to decreased CheY levels and enhanced CCW flagella rotation. B) Swimming behavior biases phagocytic uptake rates. WT flagella are mainly rotating CW, resulting in tumbling. Due to reduced numbers of cell contacts with the host cell, phagocytic uptake is low. STM Δ*fumABC* and Δ*fumABC* Δ*glgA* show both increased CCW rotation and running movement, raising the number of host cell contacts and by this phagocytic uptake. STM Δ*fumABC* Δ*glgA* exhibits prolonged CW periods, which possibly leads to slight reductions of phagocytosis rates.

## Acknowledgements

This work was supported by grants of the DFG to MH. We thank Karsten Tedin for the STM Δ*relA*/Δ*spoT* deletion strains and helpful discussions about (p)ppGpp regulation and Jürgen Heinisch (Div. Genetics) for support with qPCR.

## Materials and Methods

### Bacterial strains

*Salmonella enterica* serovar Typhimurium NCTC 12023 was used as wild-type strain (WT), and isogenic mutant strains were constructed by λ Red-mediated mutagenesis (see **Table 1**) (44). Primers and plasmids required for mutagenesis, removal of resistance cassettes and check for the correct insertion are listed in **Table 2** and **Table S 4A**. Transfer of mutant alleles into fresh strain background or for combination with other mutations occurred via P22 transduction. Both methods are described in J. Popp, et al. (45).

**Table 1.**
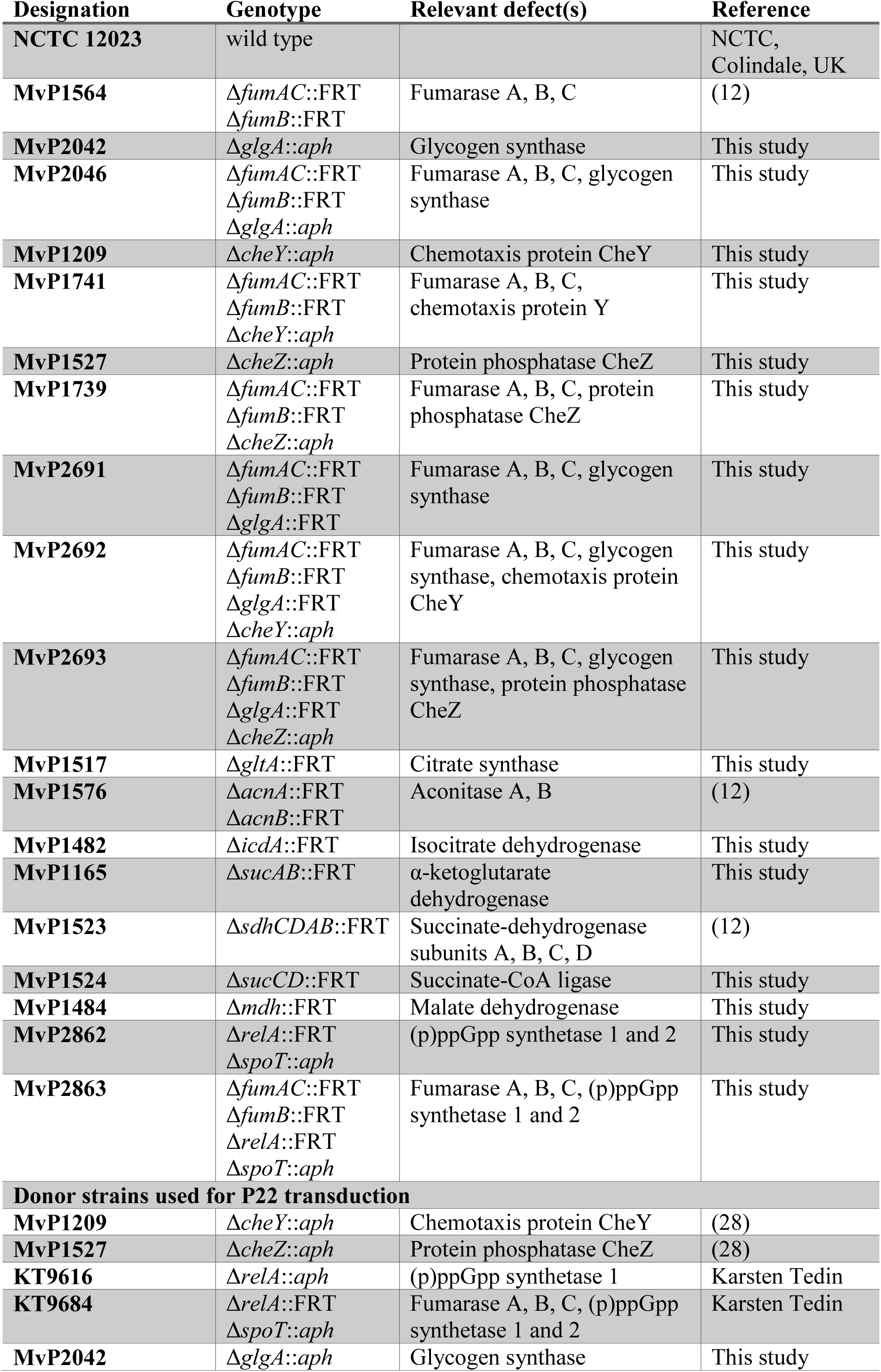
Bacterial strains used in this study.

**Table 2.**
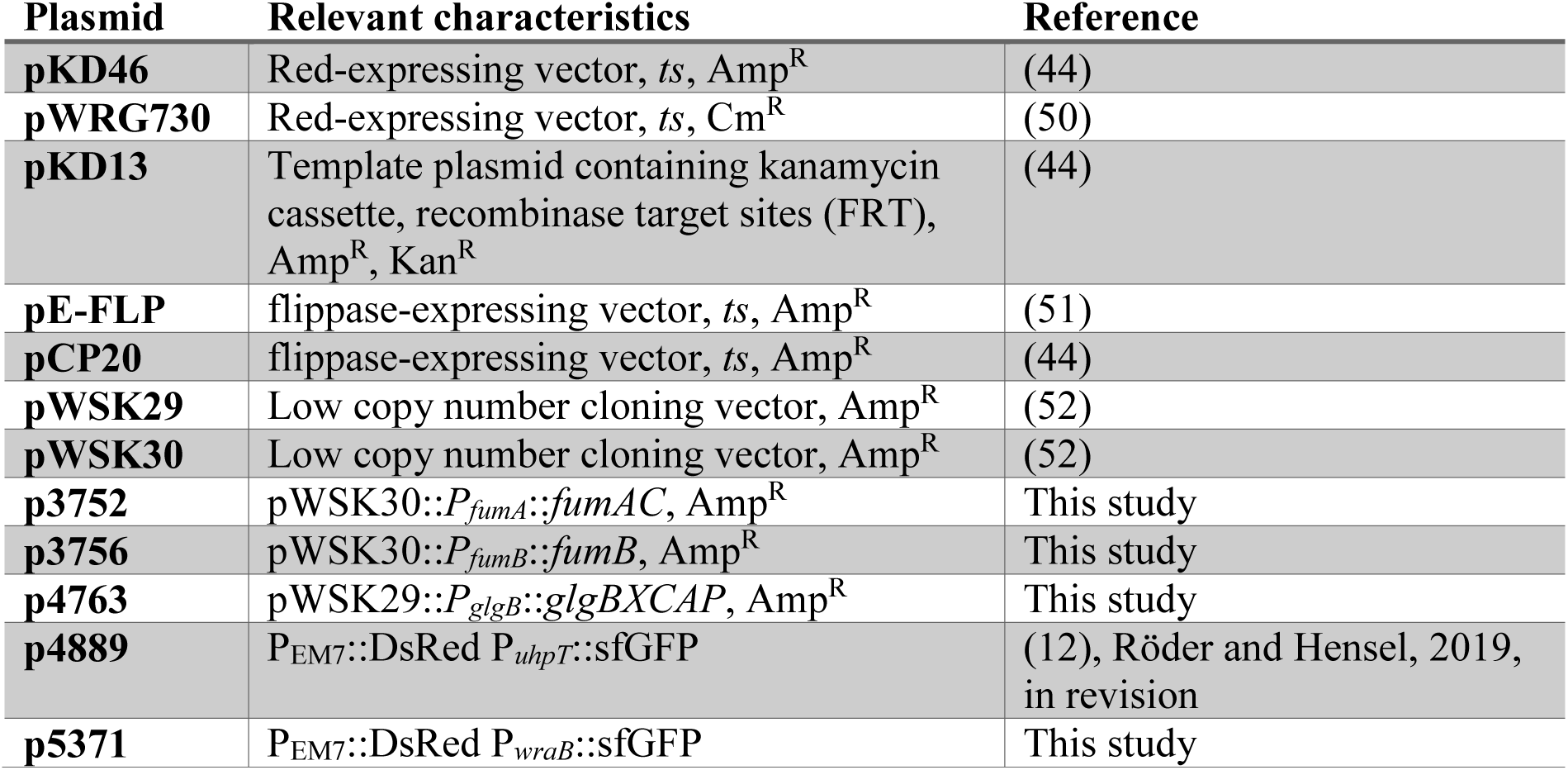
Plasmids used in this study.

### Construction of plasmids

For generation of p3752 and p3756, wild-type promoters and coding sequences of *fumAC* and *fumB* were amplified with primers listed in. After digest with NotI and XhoI or ApaI and XhoI, respectively, the gene products were ligated into the low copy plasmid pWSK30 and transformed in *E. coli* DH5α. Positive clones were confirmed with primers listed in **Table S 4A**. The plasmids were isolated and transformed in the Δ*fumAC* or Δ*fumB* deletion strain.

For construction of p4763 the promoter and sequence of *glgBXCAP* as well as the vector pWSK29 were amplified by PCR using primers listed in **Table S 4B**. The obtained PCR fragments were assembled by Gibson assembly according to the manufacturer’s protocol (NEB). Sequence-confirmed plasmids were transformed in the Δ*fumABC* Δ*glgA* deletion strain. Generation of the reporter plasmid p5330 was performed as described previously (46). Briefly, plasmid p4889 (P_EM7_::DsRed P*_uhpT_*::sfGFP) was used as vector. The *uhpT* promoter was replaced by the promoter fragment of *wraB* by Gibson assembly of fragments generated by PCR. Primers for fragment generation are listed in **Table S 4C**. Sequence-confirmed plasmids were transformed in STM WT, Δ*fumABC*, Δ*fumABC* Δ*glgA* and Δ*relA* Δ*spoT*.

### GC-MS sample preparation and measurement

Culture of strains and cell harvest occurred as described in Noster *et al.* (2019). In short: Each strain was cultured for 18.5 h at 37 °C in 25 ml LB broth with agitation at 180 rpm. For measurements of metabolites in bacterial cells, 5 ml of cultures were transferred onto Durapore PVDF filter membranes (Merck, Darmstadt, Germany) with a pore size of 0.45 µm by suction. After washing with PBS, cells were scraped from the filter into 1 ml of fresh PBS, pelleted and shock-frozen in liquid nitrogen. Afterwards samples were freeze-dried and their dry weights were determined. Metabolome analysis of the TCA cycle mutant strains was performed by GC-MS using protocols according to J. Plassmeier, et al. (47) and Noster *et al.* (2019). In short: For metabolite extraction 1 ml 80% methanol containing 10 µM ribitol (RI, internal standard) were added to dried samples and for cell disruption 500 mg acid-washed glass beads (Sigma-Aldrich, USA) and a homogenizer (Precellys, Peqlab) were used. After centrifugation, supernatants were evaporated in a nitrogen stream. For derivatization, 50 µl of a 20 mg/ml of methoxylamine hydrochloride in pyridine and N-methyl-N-[trimethylsilyl]-trifluoroacetamide were added successively to each sample and incubated with constant stirring at 37 °C for 90 min. or 30 min., respectively. RI standard was added and incubated for further 5 min. Samples were centrifuged and supernatants were used for GC-MS measurement using a TraceGC gas chromatograph equipped with a PolarisQ ion trap and an AS1000 autosampler (Thermo Finnigan, Dreieich, Germany) according to Plassmeier *et al.* (47). Metabolite quantities were normalized to ribitol and dry weights of various samples as described in Plassmeier *et al.* Mean relative pool size changes of the mutant strains compared to WT were calculated and only those data with an error probability (Student’s *t*-test) of less than 0.05 were used for further interpretation.

### Proteome profiling by LC-MS measurement

Bacteria were cultured as described for the metabolite profiling. Sample preparation and LC-MS measurement were performed according to Noster *et al*. (2019). In short, cells from 50 ml o/n culture were pelleted, suspended and washed twice with PBS. Pelleted bacteria were resuspended in lysis buffer (50 mM Tris pH 8.5, 1% SDS, protease inhibitor). Cell disruption occurred with zirconia/silica beads and a cell homogenizer. Cell debris were removed by centrifugation and proteins precipitated with TCA. Protein pellets were washed with acetone, dried and used for the following sample preparation, proteomic digest and LC-MS-measurement as described in Noster *et al*. (2019).

### Gentamicin protection assay

Culture and infection of RAW264.7 macrophages were performed as described J. Popp, et al. (45). Briefly, RAW264.7 macrophages were infected with STM o/n cultures at an MOI of 1 and centrifuged 5 min. at 370 × *g*. The infection proceeded further 25 min. Cells were washed thrice with PBS and extracellular, non-phagocytosed bacteria were killed by incubation with medium containing gentamicin (100 µg/ml for hour 1, 10 µg/ml for hour 2). 2 h p.i. cells were washed thrice with PBS and lysed by addition of 0.1% Triton-X-100 in PBS. Serial dilutions of the inoculum and lysates were plated on Mueller-Hinton II agar plates and incubated o/n at 37 °C. Phagocytosis rates were determined as percentage of internalized bacteria in dependence to the inoculum.

### Qualitative and quantitative determination of glycogen content

Qualitative determination of glycogen contents of bacterial cultures occurred as described by S. Govons, *et al*. (18). Bacterial cultures were streaked on LB agar plates and incubated o/n at 37 °C. 10 ml Lugol’s iodine solution (Roth) were added to the plate and incubated 1 min. at RT. The iodine solution was discarded and the plates photographed immediately.

Quantification of glycogen contents occurred following the protocol by N. K. Thomas Fung, Timo van der Zwan, Michael Wu (48). Of each strain, cells of 5 ml o/n culture were pelleted by centrifugation (13,000 × *g*, 10 min., 4 °C), resuspended in 50 mM TAE buffer and pelleted again. Cells were resuspended in 1.25 ml sodium acetate buffer (200 mM, pH 4.6), the suspension was added to 500 mg glass beads and disrupted by three cycles, each 1 min. with maximal speed, using Vortex Genie 2, equipped with an attachment for microtubes (scientific industries). After centrifugation, supernatants were incubated for 20 min. at 80 °C for denaturation of endogenous enzymes. For each strain 60 µl lysate were incubated with 6 µl amyloglucosidase (200 U/ml, Sigma-Aldrich) (quantification of glucose stored as glycogen and free glucose) or 6 µl water (quantification of free glucose), respectively. After incubation for 30 min. at 50 °C, 50 µl of each sample were transferred into a 96 well-plate in technical duplicates. 250 µl HK reagent (Sigma-Aldrich) were added to each sample and OD_340_ was determined in 10 min. intervals for 1 h. A standard curve with different dilutions of a glucose solution was used for extrapolation of the determined data. For relative quantification of the glycogen amount, maximal values obtained for free glucose were substracted from maximal values obtained from free glucose and glycogen and normalized to the OD_600_ of the bacterial culture.

### TEM analysis

For TEM analyses of bacteria, STM was grown o/n at 37 °C in LB broth with aeration. Cells were harvested for 2 min. at 1,250 × *g*. The pellet was resuspended in buffer (0.2 M HEPES, pH 7.4, 5 mM CaCl_2_) and bacteria were fixed by addition of glutaraldehyde (Electron Microscopy Sciences) in buffer to a final concentration of 2.5% for 1 h at 37 °C. After fixation, bacteria were washed several times in buffer and harvested for 5 min. at 625 × *g*. The pellet was gently resuspended in liquid 2% LMP agarose prewarmed to 37 °C in buffer and incubated for 10 min. at 37 °C. Bacteria in agarose were repelleted for 1 min. at 1,250 × *g* and cooled down to 4 °C until agarose was solid. The agarose block containing the bacteria was cut into small cubes (max. 1 mm^3^) and cubes were post-fixed was performed with 2% osmium tetroxide (Electron Microscopy Sciences) in buffer containing 1.5% potassium ferricyanide (Sigma) and 0.1% ruthenium red (Applichem) for 1.5 h at 4 °C in the dark. After several washing steps, bacteria were dehydrated in a cold graded ethanol series and finally rinsed in anhydrous ethanol at RT twice. The agarose cubes were flat-embedded in EPON812 (Serva). Serial 70 nm sections were generated with an ultramicrotome (Leica EM UC6) and collected on formvar-coated EM copper grids. After staining with uranyl acetate and lead citrate, bacteria were observed with TEM (Zeiss EFTEM 902 A), operated at 80 kV and equipped with a 2K wide-angle slow-scan CCD camera (TRS, Moorenwies, Germany). Images were taken with the software ImageSP (TRS image SysProg, Moorenwies, Germany).

### Flagella rotation analysis

Flagella rotation was determined as illustrated in **Fig. S 5**. Bacteria were cultured for 18.5 h in LB, diluted 1:100 in PBS and subjected to shear force by pressing the suspension eight times through a syringe equipped with a 24 G cannula. 15 µl sample were placed onto a microscope slide and covered with a polystyrene-coated coverslip, on which three small drops of vacuum grease were spotted to achieve an optimal distance allowing free movement of STM. Sealing the cover slip with a 1:1:1 mixture of vaseline, lanoline and paraffin avoided suction. Rotating cells, bound with their flagella filaments to the coverslip, were selected and rotation direction was recorded using the Axio Observer microscope with an AxioCam CCD camera (Zeiss) for periods of 18 s (frame rate 57/s). After image processing with Fiji (size reduction, background subtraction, contrast enhancement, smoothing with the GaussianBlur plug-in filter), rotation analyses were performed using the tool SpinningBug Tracker (user-written software, Matlab 7.17 (R2012a)). By detection of the angle between the rotating bacteria and a reference axis, the rotation direction was calculated. CCW rotation of the bacterial body has to be interpreted as CW rotation of the flagellum and *vice versa*. Rotations of less than 5° per frame were defined as pause. Bacteria rotating with speeds of > 180°/frame were excluded, due to limited time-resolution. Switching events were defined as changes from CW to CCW rotation and *vice versa*.

### Swimming path analysis

Bacteria were cultured 18.5 h with aeration in LB and diluted 1:20 in PBS. The assembly of microcopy slide, sample and coverslip was similar as described for the flagella rotation analysis, but without prior coating of the coverslip with polystyrene. The swimming bacteria were recorded for 100 frames (14 frames/s). Visualization of swimming paths was performed with ImageJ, using the plug-in MTrackJ (49).

### qPCR

For RNA preparation by ‘hot phenol’ method, bacteria were cultured 18.5 h in LB with aeration. 1.2 × 10^9^ bacteria were pelleted, treated with stop-solution (95% EtOH, 5% phenol saturated with 0.1 M citrate buffer, pH 4.3 (Sigma-Aldrich) and snap-frozen in liquid nitrogen. All following steps were conducted as described in detail in Noster *et al.* (2019) according to protocols from Mattatall and Sanderson (1996) and Sittka *et al.* (2009). For cDNA synthesis the RevertAid First strand cDNA synthesis kit (ThermoFisher Scientific) was used, applying 1 µg RNA and random hexamer primers. qPCR was performed using the Maxima SYBR Green/Fluorescein qPCR Master Mix (ThermoFisher) and iCycler equipped with MyiQ module (Biorad). Data were normalized to expression levels of a house-keeping gene (16S rRNA) and calculated in consideration of primer efficiencies, which were determined before using serial dilutions of cDNA. Oligonucleotides used are listed in **Table S 4D**.

### Flow cytometry analyses

STM strains harboring p5330 were grown in LB broth at 37 °C with aeration for 18.5 h, diluted 1:1000 in FACS buffer (1 % BSA in PBS, 1 mM EDTA, 20 mM Hepes pH 7.2, 50 mM NH_4_Cl) and subjected to flow cytometry on an Attune NxT instrument (Thermo Fischer Scientific). The intensity of the sfGFP fluorescence per gated STM cell of 10,000 bacteria with constitutive red fluorescence was recorded and x-medians for sfGFP intensities were calculated.

## Suppl. Materials

### Suppl. Tables

**Table S 1.** Metabolomics, TCA cycle enzyme deletion strains, *glgA*, *fumABC glgA* strains

**Table S 2.** Proteomics, WT vs. *fumABC* strains

**Table S 3.** Proteomics, *fumABC* vs. *fumABC glgA* strains

**Table S 4.**
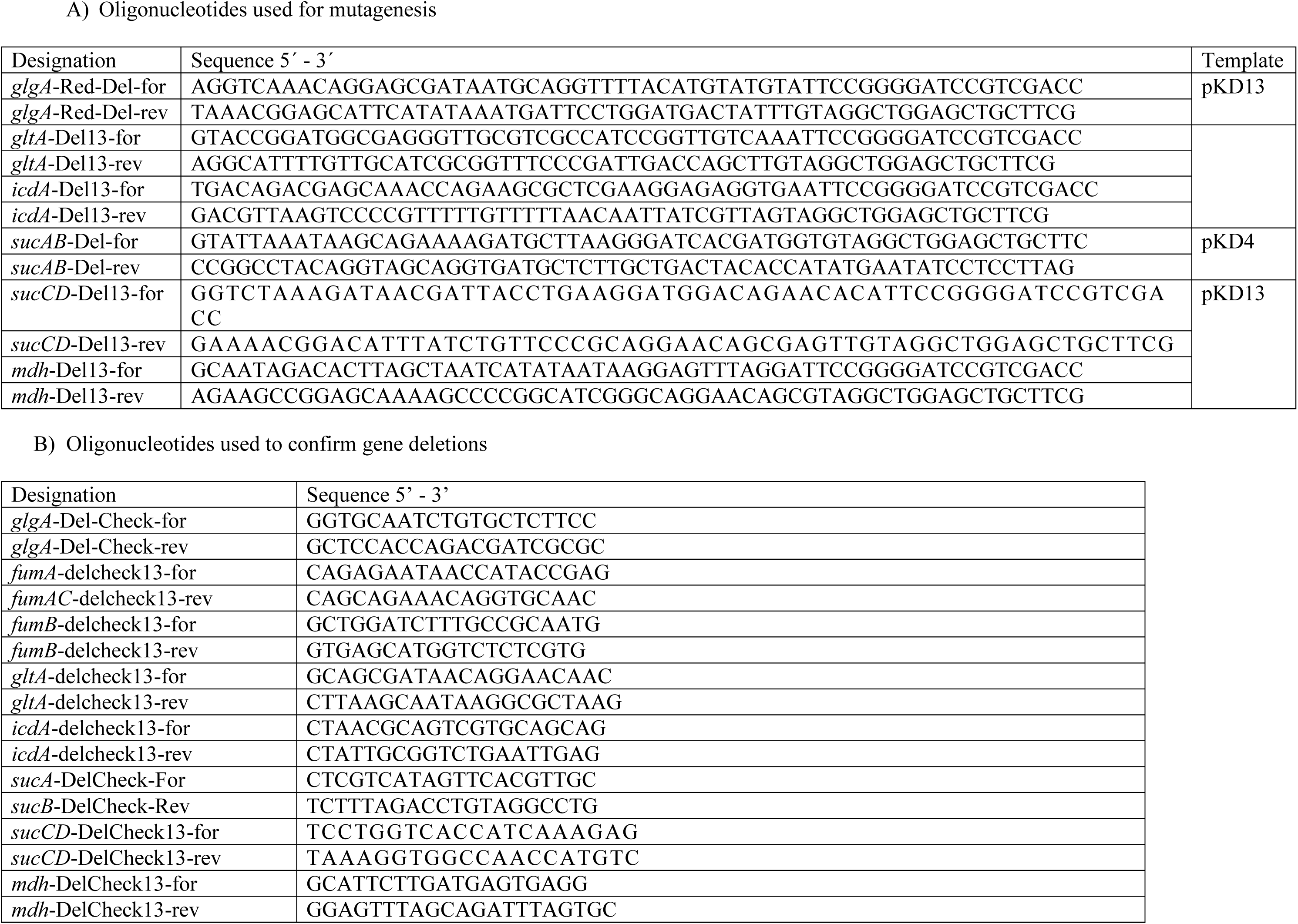
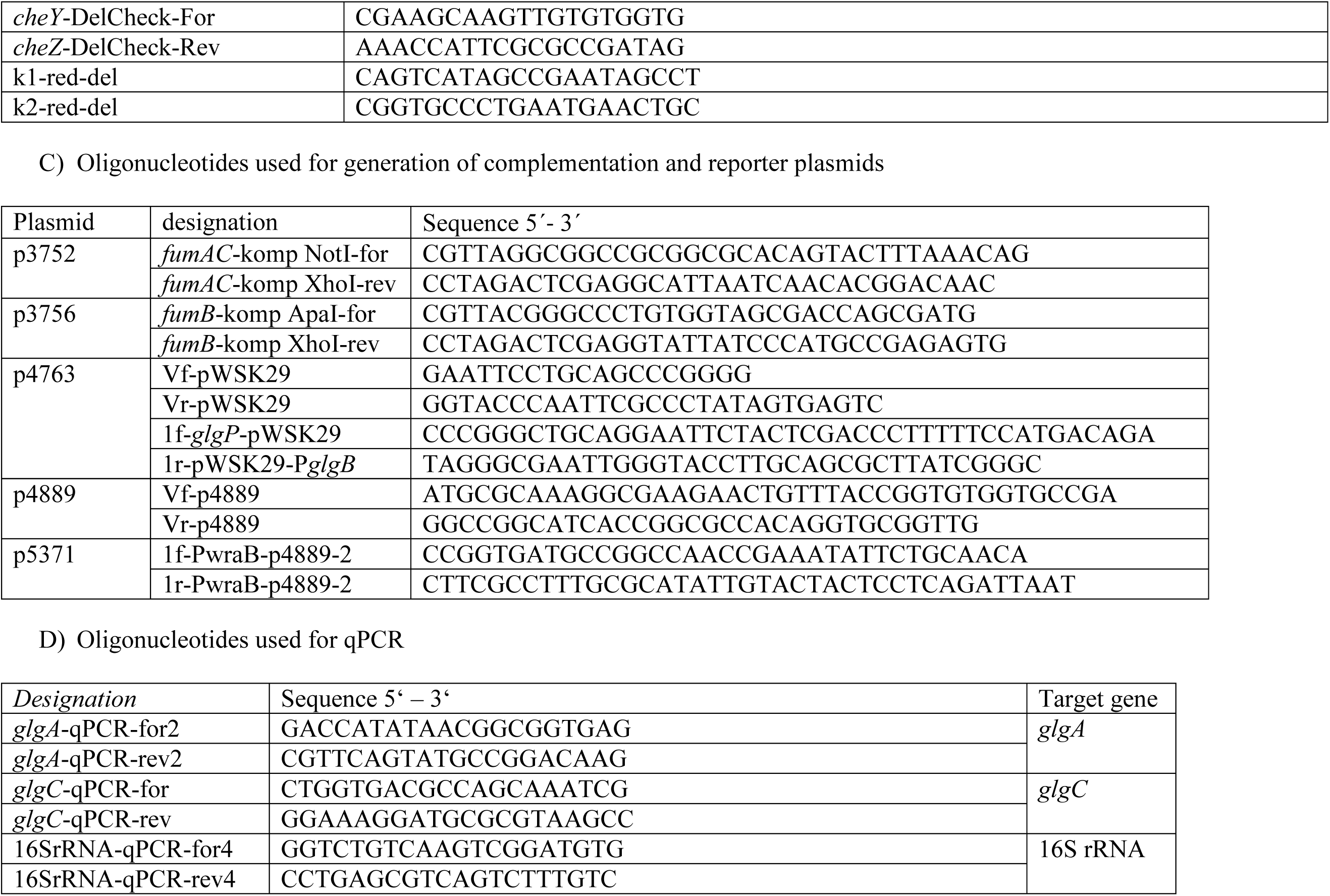
Oligonucleotides used in this study

### Suppl. Figures and Legends

**Fig. S 1.**
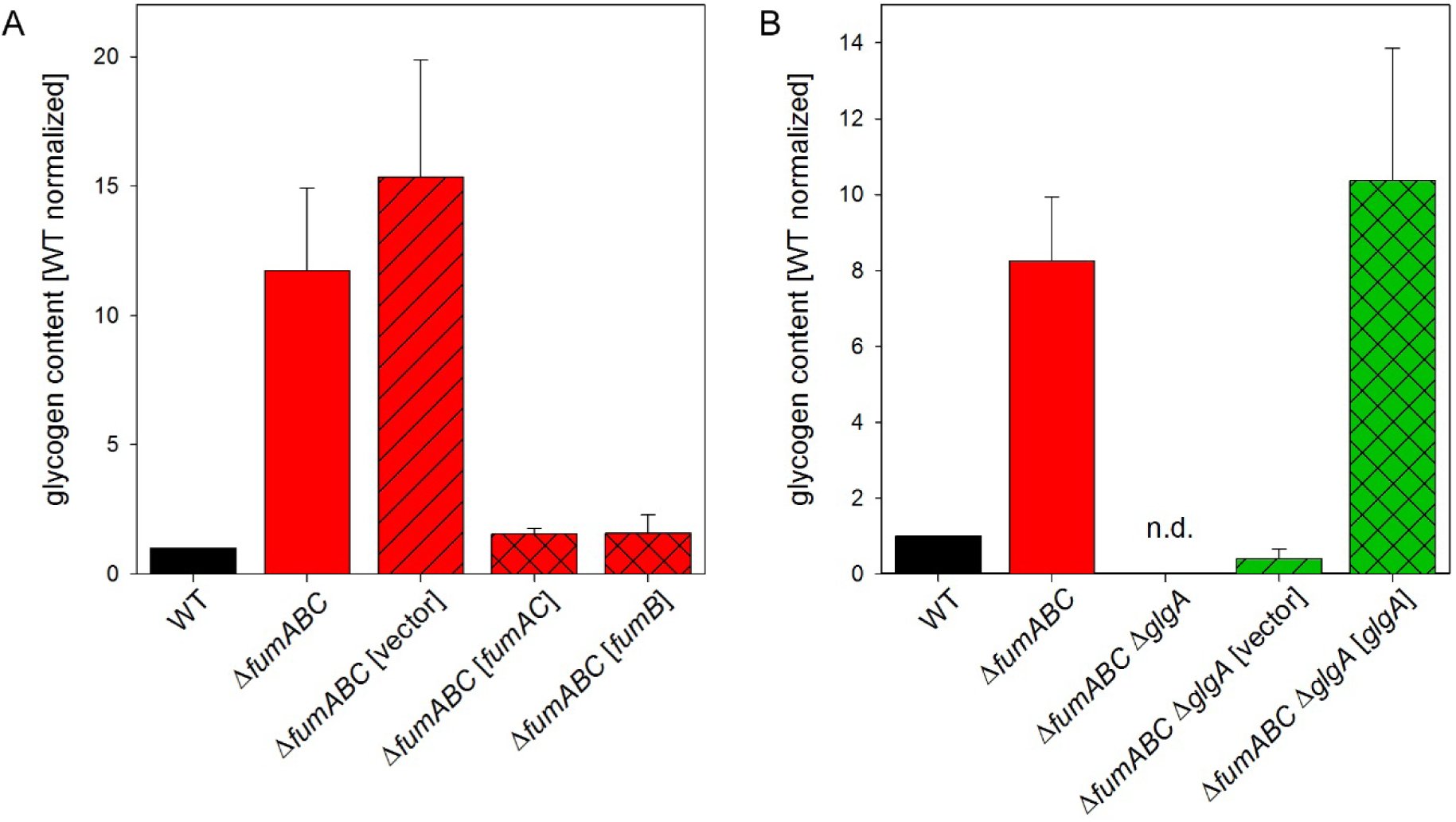
Glycogen accumulation in the Δ*fumABC* mutant strain is dependent on the deletion of all fumarase isoforms and an intact glycogen synthase. Quantification of glycogen contents of STM strains occurred as described in Fig. 3. A) Complementation of the Δ*fumABC* deletion strain harboring plasmids encoding the intact genes *fumAC* or *fumB* gene under control of their native promoter or the empty vector, respectively. B) Complementation of the Δ*fumABC* Δ*glgA* deletion strain with plasmids encoding the intact gene *glgA* or the empty vector, respectively. Glycogen concentrations were normalized to WT (=1), error bars represent standard deviations of two independent biological replicates, each consisting of two technical replicates.

**Fig. S 2.**
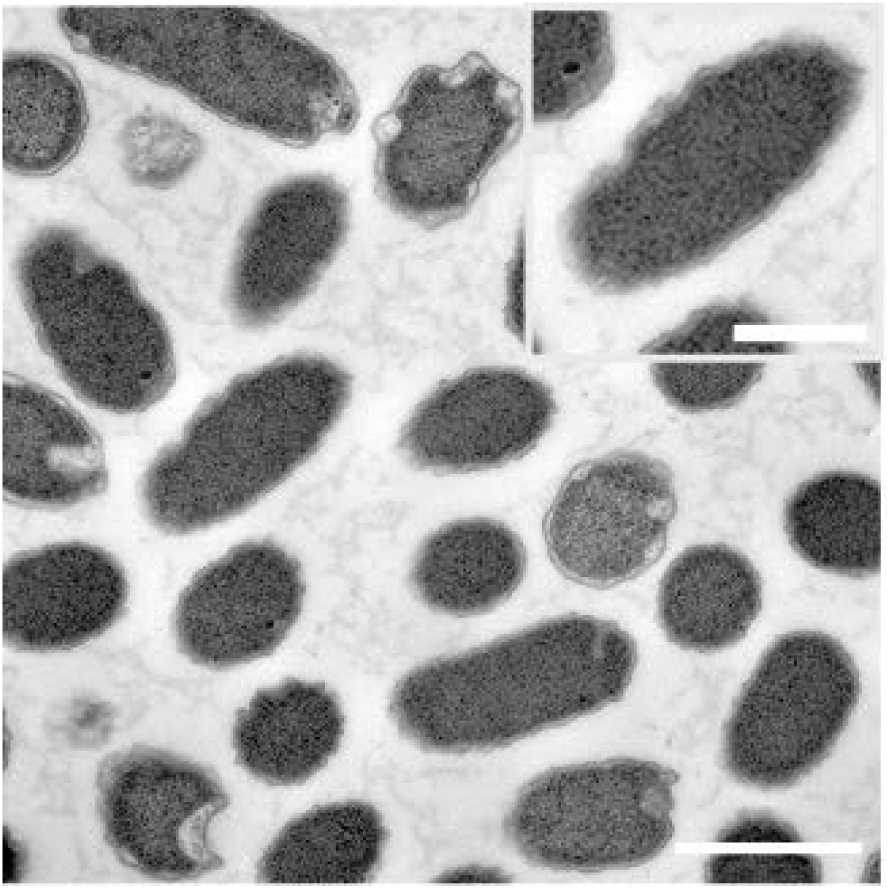
Absence of granula accunulation in STM Δ*fumABC* by glycogen synthase deletion. Electron micrographs of STM Δ*fumABC* Δ*glgA*, aerobically grown for 18.5 h in LB broth. Note the absence of polymers accumulations of in the polar regions of the bacterium. Scale bars, 1 µm (overview), 0.5 µm (detail).

**Fig. S 3.**
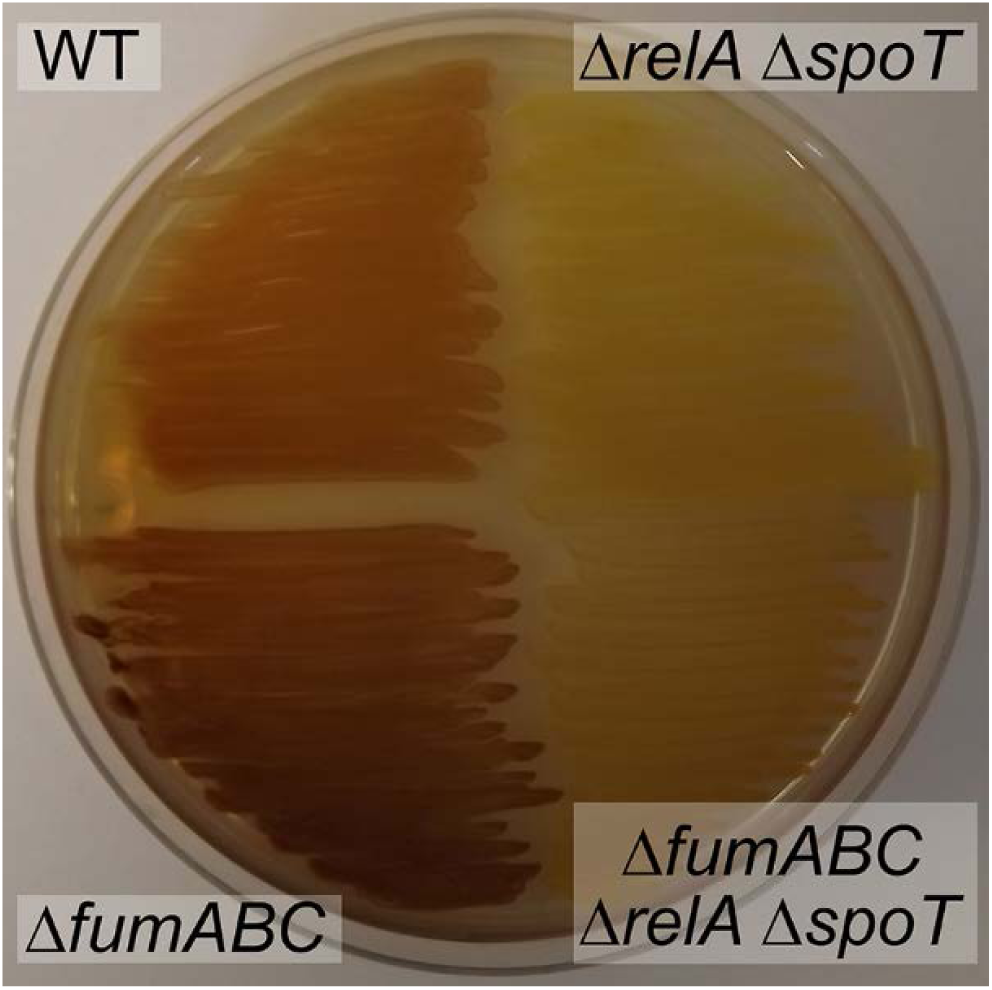
Glycogen accumulation in Δ*fumABC* depends on (p)ppGpp synthesizing enzymes. STM WT, Δ*fumABC*, Δ*relA spoT* and Δ*fumABC* Δ*relA* Δ*spoT* strains were grown on LB agar plates for 18.5 h at 37 °C. Potassium iodine staining was performed and brownish color indicates intercalations of iodine with glycogen.

**Fig. S 4.**
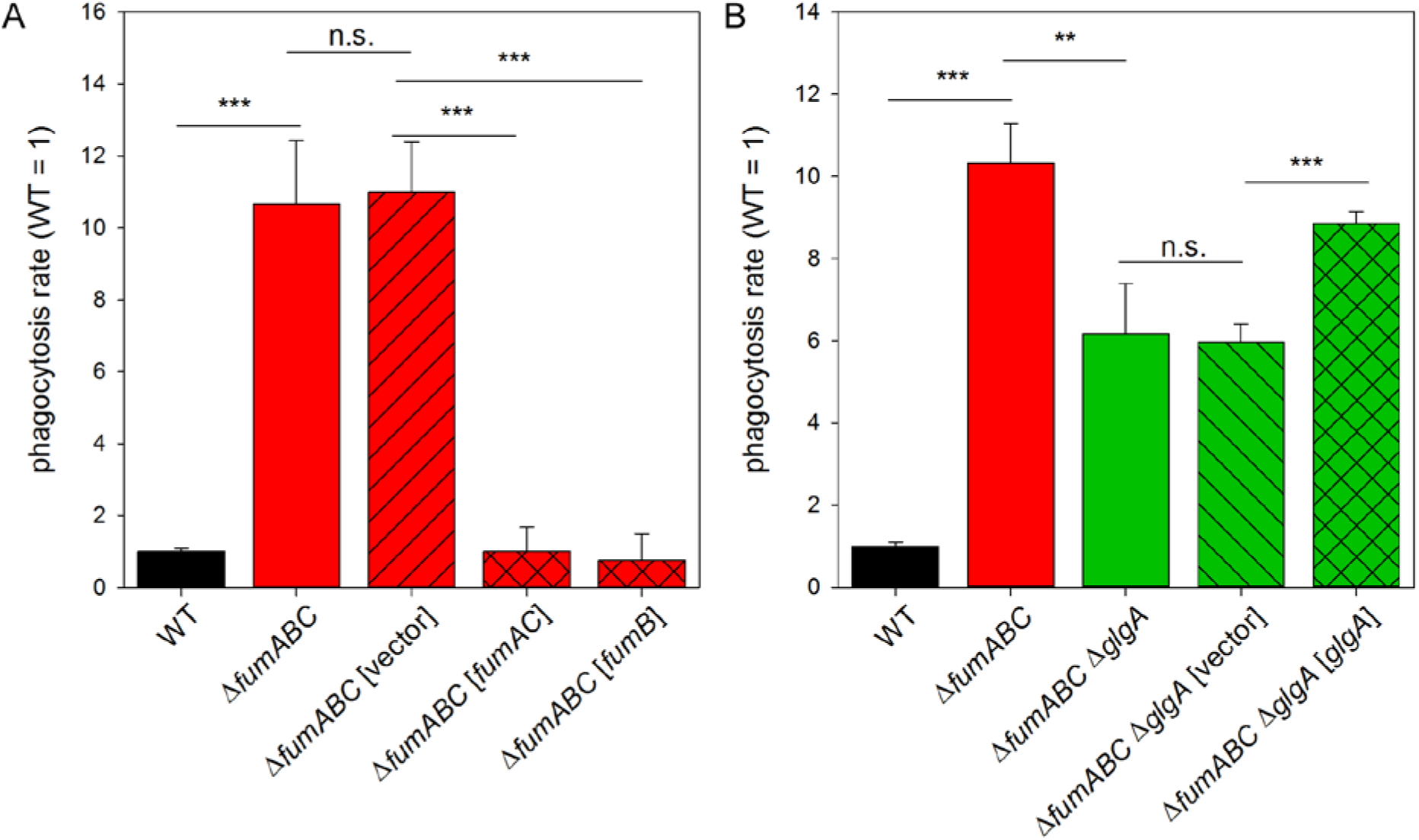
Deletion of fumarases increases the phagocytic uptake by RAW264.7 macrophages and is dependent on glycogen synthesis. RAW264.7 macrophages were infected as described for Fig. 8. Values were normalized to WT (=1), and means and standard deviations of three technical replicates are shown. A) RAW264.7 macrophages were infected with WT, Δ*fumABC* and Δ*fumABC* strains harboring plasmids encoding the intact genes *fumAC* or *fumB* gene under control of their native promoter or the empty vector, respectively. B) RAW264.7 macrophages were infected with WT, Δ*fumABC*, Δ*fumABC* Δ*glgA* and Δ*fumABC* Δ*glgA* strains harboring a plasmid encoding wild-type GlgA under control of its native promoter, or the empty vector, respectively. The data are representative for three independent biological replicates. Statistical analyses were performed by Student’s *t*-test and significances are indicated as follows: **, *p* < 0.01; ***, *p* < 0.001; n.s., not significant.

**Fig. S 5.**
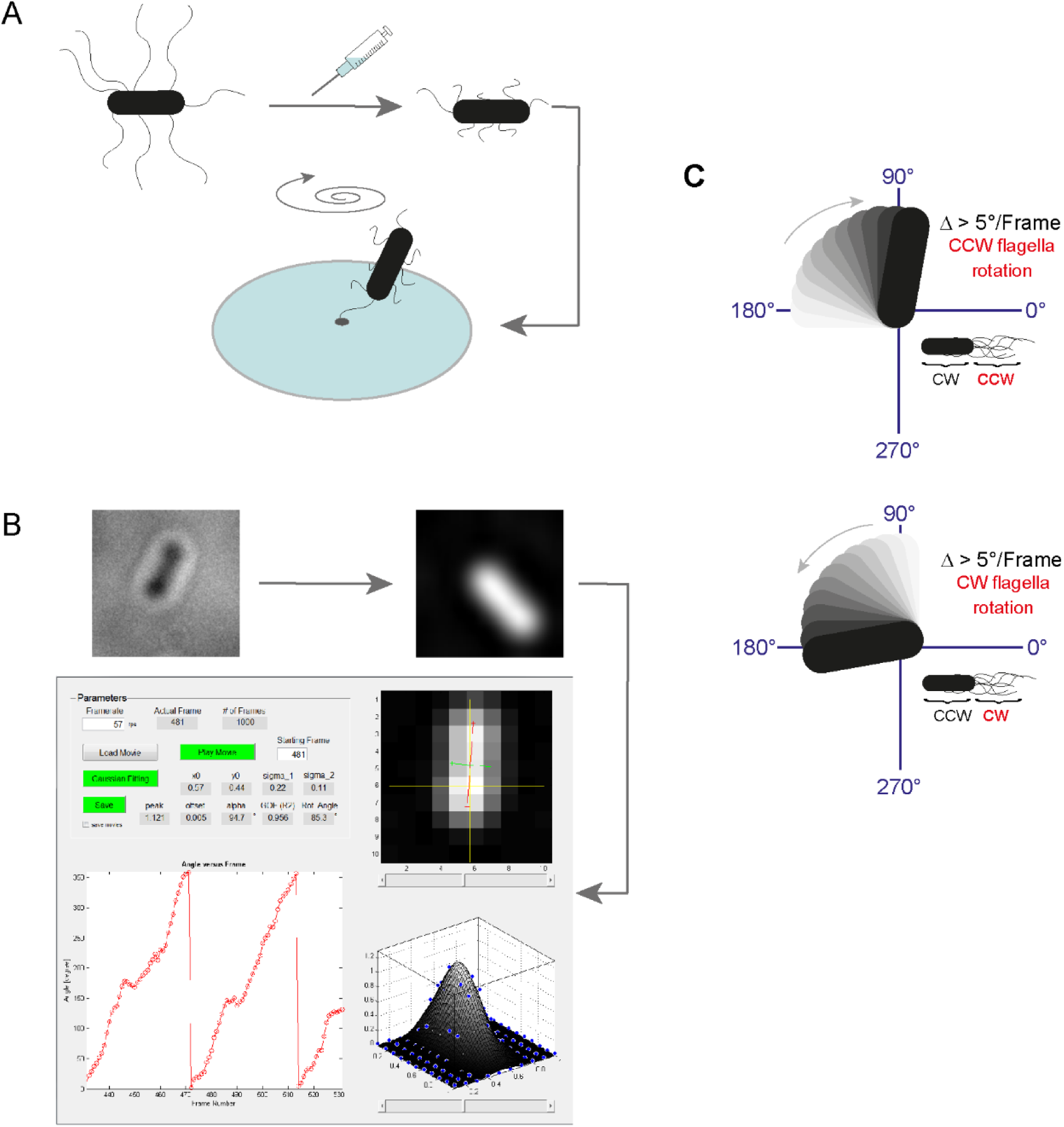
Method of flagella rotation analysis. Bacteria were cultured for 18.5 h in LB and diluted 1:100 in PBS. A) Bacteria were subjected to shearing to remove and shorten flagellar filaments. A small volume was given on an object slide and covered with a polystyrene-coated coverslip. Distance between object slide and coverslip was achieved by drops of vacuum grease. Dehydration or undertow were avoided by sealing the coverslip with *valap* (1:1:1 mixture of vaselin, lanolin and paraffin). Bacteria fixed with only one flagellum to the coverslip showed rotation of the body, which was recorded using the Axio Observer microscope with an AxioCam CCD camera (Zeiss) for periods of 17.54 s (frame rate: 57 frames/s). B) Videos of rotating bacteria were processed using ImageJ and rotation analyses were performed using the tool SpinningBug Tracker. By detection of the angle between the rotating bacteria and a reference axis, the rotation direction was calculated. C) If there was a change of degree of more than 5° it was defined as rotation. The visible clockwise rotation of the bacterial body results from counterclockwise rotation of a flagellum, and *vice versa*.

## References

1. Vuoristo KS, Mars AE, Sanders JPM, Eggink G, Weusthuis RA. 2016. Metabolic Engineering of TCA Cycle for Production of Chemicals. Trends Biotechnol 34:191–197.

2. Durica-Mitic S, Gopel Y, Gorke B. 2018. Carbohydrate Utilization in Bacteria: Making the Most Out of Sugars with the Help of Small Regulatory RNAs. Microbiol Spectr 6.

3. Nimmo GA, Nimmo HG. 1984. The regulatory properties of isocitrate dehydrogenase kinase and isocitrate dehydrogenase phosphatase from Escherichia coli ML308 and the roles of these activities in the control of isocitrate dehydrogenase. Eur J Biochem 141:409–414.

4. Tretter L, Adam-Vizi V. 2005. Alpha-ketoglutarate dehydrogenase: a target and generator of oxidative stress. Philos Trans R Soc Lond B Biol Sci 360:2335–2345.

5. Weitzman PD. 1966. Regulation of citrate synthase activity in escherichia coli. Biochim Biophys Acta 128:213–215.

6. Gu M, Imlay JA. 2011. The SoxRS response of *Escherichia coli* is directly activated by redox-cycling drugs rather than by superoxide. Mol Microbiol 79:1136–1150.

7. Kohanski MA, Dwyer DJ, Hayete B, Lawrence CA, Collins JJ. 2007. A common mechanism of cellular death induced by bactericidal antibiotics. Cell 130:797–810.

8. Cameron EA, Sperandio V. 2015. Frenemies: Signaling and Nutritional Integration in Pathogen-Microbiota-Host Interactions. Cell Host Microbe 18:275–284.

9. Deiwick J, Nikolaus T, Erdogan S, Hensel M. 1999. Environmental regulation of Salmonella pathogenicity island 2 gene expression. Mol Microbiol 31:1759–1773.

10. van der Heijden J, Bosman ES, Reynolds LA, Finlay BB. 2015. Direct measurement of oxidative and nitrosative stress dynamics in Salmonella inside macrophages. Proc Natl Acad Sci U S A 112:560–565.

11. Becker D, Selbach M, Rollenhagen C, Ballmaier M, Meyer TF, Mann M, Bumann D. 2006. Robust Salmonella metabolism limits possibilities for new antimicrobials. Nature 440:303–307.

12. Noster J, Persicke M, Chao TC, Krone L, Heppner B, Hensel M, Hansmeier N. 2019. Impact of ROS-Induced Damage of TCA Cycle Enzymes on Metabolism and Virulence of Salmonella enterica serovar Typhimurium. Front Microbiol 10:762.

13. Kuo CJ, Wang ST, Lin CM, Chiu HC, Huang CR, Lee DY, Chang GD, Chou TC, Chen JW, Chen CS. 2018. A multi-omic analysis reveals the role of fumarate in regulating the virulence of enterohemorrhagic Escherichia coli. Cell Death Dis 9:381.

14. Ruecker N, Jansen R, Trujillo C, Puckett S, Jayachandran P, Piroli GG, Frizzell N, Molina H, Rhee KY, Ehrt S. 2017. Fumarase Deficiency Causes Protein and Metabolite Succination and Intoxicates Mycobacterium tuberculosis. Cell Chem Biol 24:306–315.

15. Kim JS, Cho DH, Heo P, Jung SC, Park M, Oh EJ, Sung J, Kim PJ, Lee SC, Lee DH, Lee S, Lee CH, Shin D, Jin YS, Kweon DH. 2016. Fumarate-Mediated Persistence of Escherichia coli against Antibiotics. Antimicrob Agents Chemother 60:2232–2240.

16. Barak R, Eisenbach M. 1992. Fumarate or a fumarate metabolite restores switching ability to rotating flagella of bacterial envelopes. J Bacteriol 174:643–645.

17. Prasad K, Caplan SR, Eisenbach M. 1998. Fumarate modulates bacterial flagellar rotation by lowering the free energy difference between the clockwise and counterclockwise states of the motor. J Mol Biol 280:821–828.

18. Govons S, Vinopal R, Ingraham J, Preiss J. 1969. Isolation of mutants of Escherichia coli B altered in their ability to synthesize glycogen. J Bacteriol 97:970–972.

19. Preiss J. 2009. Glycogen: Biosynthesis and Regulation. EcoSal Plus 3.

20. Moran-Zorzano MT, Alonso-Casajus N, Munoz FJ, Viale AM, Baroja-Fernandez E, Eydallin G, Pozueta-Romero J. 2007. Occurrence of more than one important source of ADPglucose linked to glycogen biosynthesis in Escherichia coli and Salmonella. FEBS Lett 581:4423–4429.

21. Eydallin G, Montero M, Almagro G, Sesma MT, Viale AM, Munoz FJ, Rahimpour M, Baroja-Fernandez E, Pozueta-Romero J. 2010. Genome-wide screening of genes whose enhanced expression affects glycogen accumulation in Escherichia coli. DNA Res 17:61–71.

22. Amato SM, Orman MA, Brynildsen MP. 2013. Metabolic control of persister formation in Escherichia coli. Mol Cell 50:475–487.

23. Tedin K, Norel F. 2001. Comparison of DeltarelA strains of Escherichia coli and Salmonella enterica serovar Typhimurium suggests a role for ppGpp in attenuation regulation of branched-chain amino acid biosynthesis. J Bacteriol 183:6184–6196.

24. Rahimpour M, Montero M, Almagro G, Viale AM, Sevilla A, Canovas M, Munoz FJ, Baroja-Fernandez E, Bahaji A, Eydallin G, Dose H, Takeuchi R, Mori H, Pozueta-Romero J. 2013. GlgS, described previously as a glycogen synthesis control protein, negatively regulates motility and biofilm formation in Escherichia coli. Biochem J 452:559–573.

25. Montrone M, Eisenbach M, Oesterhelt D, Marwan W. 1998. Regulation of switching frequency and bias of the bacterial flagellar motor by CheY and fumarate. J Bacteriol 180:3375–3380.

26. Wadhams GH, Armitage JP. 2004. Making sense of it all: bacterial chemotaxis. Nat Rev Mol Cell Biol 5:1024–1037.

27. Silverman M, Simon M. 1974. Flagellar rotation and the mechanism of bacterial motility. Nature 249:73–74.

28. Lorkowski M, Felipe-Lopez A, Danzer CA, Hansmeier N, Hensel M. 2014. Salmonella enterica invasion of polarized epithelial cells is a highly cooperative effort. Infect Immun 82:2657–2667.

29. Misselwitz B, Barrett N, Kreibich S, Vonaesch P, Andritschke D, Rout S, Weidner K, Sormaz M, Songhet P, Horvath P, Chabria M, Vogel V, Spori DM, Jenny P, Hardt WD. 2012. Near surface swimming of Salmonella Typhimurium explains target-site selection and cooperative invasion. PLoS Pathog 8:e1002810.

30. Achouri S, Wright JA, Evans L, Macleod C, Fraser G, Cicuta P, Bryant CE. 2015. The frequency and duration of Salmonella-macrophage adhesion events determines infection efficiency. Philos Trans R Soc Lond B Biol Sci 370:20140033.

31. Jones BD, Lee CA, Falkow S. 1992. Invasion by Salmonella typhimurium is affected by the direction of flagellar rotation. Infect Immun 60:2475–2480.

32. Parkinson JS. 1978. Complementation analysis and deletion mapping of Escherichia coli mutants defective in chemotaxis. J Bacteriol 135:45–53.

33. Morin M, Ropers D, Letisse F, Laguerre S, Portais JC, Cocaign-Bousquet M, Enjalbert B. 2016. The post-transcriptional regulatory system CSR controls the balance of metabolic pools in upper glycolysis of Escherichia coli. Mol Microbiol 100:686–700.

34. Revelles O, Millard P, Nougayrede JP, Dobrindt U, Oswald E, Letisse F, Portais JC. 2013. The carbon storage regulator (Csr) system exerts a nutrient-specific control over central metabolism in Escherichia coli strain Nissle 1917. PLoS One 8:e66386.

35. Pannuri A, Yakhnin H, Vakulskas CA, Edwards AN, Babitzke P, Romeo T. 2012. Translational repression of NhaR, a novel pathway for multi-tier regulation of biofilm circuitry by CsrA. J Bacteriol 194:79–89.

36. Sterzenbach T, Nguyen KT, Nuccio SP, Winter MG, Vakulskas CA, Clegg S, Romeo T, Baumler AJ. 2013. A novel CsrA titration mechanism regulates fimbrial gene expression in Salmonella typhimurium. EMBO J 32:2872–2883.

37. Montero M, Rahimpour M, Viale AM, Almagro G, Eydallin G, Sevilla A, Canovas M, Bernal C, Lozano AB, Munoz FJ, Baroja-Fernandez E, Bahaji A, Mori H, Codoner FM, Pozueta-Romero J. 2014. Systematic production of inactivating and non-inactivating suppressor mutations at the relA locus that compensate the detrimental effects of complete spot loss and affect glycogen content in Escherichia coli. PLoS One 9:e106938.

38. Amato SM, Brynildsen MP. 2015. Persister Heterogeneity Arising from a Single Metabolic Stress. Curr Biol 25:2090–2098.

39. Traxler MF, Zacharia VM, Marquardt S, Summers SM, Nguyen HT, Stark SE, Conway T. 2011. Discretely calibrated regulatory loops controlled by ppGpp partition gene induction across the ‘feast to famine’ gradient in Escherichia coli. Mol Microbiol 79:830–845.

40. Lazzarini RA, Cashel M, Gallant J. 1971. On the regulation of guanosine tetraphosphate levels in stringent and relaxed strains of Escherichia coli. J Biol Chem 246:4381–4385.

41. Fibriansah G, Veetil VP, Poelarends GJ, Thunnissen AM. 2011. Structural basis for the catalytic mechanism of aspartate ammonia lyase. Biochemistry 50:6053–6062.

42. Kanehisa M, Goto S. 2000. KEGG: kyoto encyclopedia of genes and genomes. Nucleic Acids Res 28:27–30.

43. Tran QH, Unden G. 1998. Changes in the proton potential and the cellular energetics of Escherichia coli during growth by aerobic and anaerobic respiration or by fermentation. Eur J Biochem 251:538–543.

44. Datsenko KA, Wanner BL. 2000. One-step inactivation of chromosomal genes in Escherichia coli K-12 using PCR products. Proc Natl Acad Sci U S A 97:6640–6645.

45. Popp J, Noster J, Busch K, Kehl A, Zur Hellen G, Hensel M. 2015. Role of host cell-derived amino acids in nutrition of intracellular Salmonella enterica. Infect Immun 83:4466–4475.

46. Noster J, Chao TC, Sander N, Schulte M, Reuter T, Hansmeier N, Hensel M. 2019. Proteomics of intracellular Salmonella enterica reveals roles of Salmonella pathogenicity island 2 in metabolism and antioxidant defense. PLoS Pathog 15:e1007741.

47. Plassmeier J, Barsch A, Persicke M, Niehaus K, Kalinowski J. 2007. Investigation of central carbon metabolism and the 2-methylcitrate cycle in Corynebacterium glutamicum by metabolic profiling using gas chromatography-mass spectrometry. J Biotechnol 130:354–363.

48. Thomas Fung NK, Timo van der Zwan, Michael Wu. 2013. Residual glycogen metabolism in Escherichia coli is specific to the limiting macronutrient and varies during stationary phase. Journal of Experimental Microbiology and Immunology 17:83–87.

49. Meijering E, Dzyubachyk O, Smal I. 2012. Methods for cell and particle tracking. Methods Enzymol 504:183–200.

50. Hoffmann S, Schmidt C, Walter S, Bender JK, Gerlach RG. 2017. Scarless deletion of up to seven methyl-accepting chemotaxis genes with an optimized method highlights key function of CheM in Salmonella Typhimurium. PLoS One 12:e0172630.

51. St-Pierre F, Cui L, Priest DG, Endy D, Dodd IB, Shearwin KE. 2013. One-step cloning and chromosomal integration of DNA. ACS Synth Biol 2:537–541.

52. Wang RF, Kushner SR. 1991. Construction of versatile low-copy-number vectors for cloning, sequencing and gene expression in Escherichia coli. Gene 100:195–199.

